# Gut environmental factors explain variations in the gut microbiome composition and metabolism within and between healthy adults

**DOI:** 10.1101/2024.01.23.574598

**Authors:** Nicola Procházková, Martin F. Laursen, Giorgia La Barbera, Eirini Tsekitsidi, Malte S. Jørgensen, Morten A. Rasmussen, Jeroen Raes, Tine R. Licht, Lars O. Dragsted, Henrik M. Roager

## Abstract

The human gut microbiome is highly personal. However, the contribution of the gut environment to variations in the gut microbiome remains elusive. Here, we profiled the gut microbiome composition and metabolism over 9 consecutive days in 61 healthy adults and assessed gut environmental factors including segmental transit time and pH using a wireless motility capsule. Day-to-day fluctuations in gut environmental factors as well as segmental transit time and pH varied substantially between individuals. The gut environment explained more variations in gut microbiome and urine metabolome than dietary macronutrients or personal characteristics. Finally, we identified coffee-derived metabolites to be negatively correlated with small intestinal transit time and several microbial metabolites to be associated with colonic transit time including urinary proteolytic markers, faecal short-chain fatty acids, and breath methane. Our work suggests that the gut environment is key for understanding the individuality of the human gut microbiome composition and metabolism.

## Introduction

Diet provides substrates to the residents of the human gut and thereby influences the microbial composition and metabolism^1,2^. However, inter-individual variation in the gut microbiome composition is observed even with identical dietary intake^3^, and not all microbial-derived metabolites are equally sensitive to dietary changes^4^, suggesting that other factors in the gut contribute to the variations in microbial metabolism. Gut transit time accounts for a large proportion of both inter- and intra-individual variation in the microbiome composition of healthy populations^5–8^. Long transit time through the whole gastrointestinal tract (GIT) is associated with changes in microbial metabolism towards increased protein degradation and methane production^9^. While the main microbial products of saccharolysis, i.e. short-chain fatty acids (SCFAs), are typically considered beneficial for the host^10^, microbial proteolysis results in metabolites associated with poor health outcomes, including as hydrogen sulfide, ammonia, branched-chain fatty acids (BCFAs), p-cresol, indole, and phenylacetate^11,12^. The marked changes in pH along the GIT are also linked to gut microbial composition and metabolism^13^. For instance, the presence of SCFAs and other organic acids such as lactate or succinate produced by the gut microbiota lowers the colonic pH^13^, which in turn inhibits bacteria sensitive to acidic environments^14^. Nonetheless, little is known about how the gut environmental factors, such as segmental transit times and GIT luminal pH variation, associate with diet-host-microbiota metabolism and thereby account for differences within and between healthy adults. Understanding how these factors associate with the host-microbiota metabolism could be crucial for developing future personalized dietary microbiome-based strategies. We, therefore, conducted a 9-day human study including 61 healthy volunteers (PRIMA, ClinicalTrials.gov identifier: NCT04804319), residing in Denmark. We combined assessment of whole gut and segmental gastrointestinal transit time and pH with bowel habits (i.e. stool consistency, stool frequency, and stool moisture), 24-h dietary records, measurements of breath hydrogen and methane, and multi-omics profiling of urine and faecal samples. The longitudinal study design and repeated sampling allowed us to follow inter-individual and day-to-day changes in the gut environmental factors, gut microbiota, and microbiota-derived metabolites including levels of faecal SCFAs and BCFAs, as well as urinary levels of products of microbial proteolysis.

## Results

### Study design and participants’ characteristics

Our study was designed to explore links between gut environmental factors and variations in gut microbiome composition and host-microbiota co-metabolism within and between individuals (**Figure 1A**). To characterize these links, 61 healthy participants were enrolled (age 39 ± 13.5 years, BMI 23.6 ± 2.8 kg/m^2^, see **Table 1** for participants’ characteristics and **Methods** for enrollment criteria) and asked to maintain their habitual lifestyle and diet for 9 consecutive days. All enrolled participants completed the 9-days trial (**Supplementary Fig. 1**)

**Figure 1.**
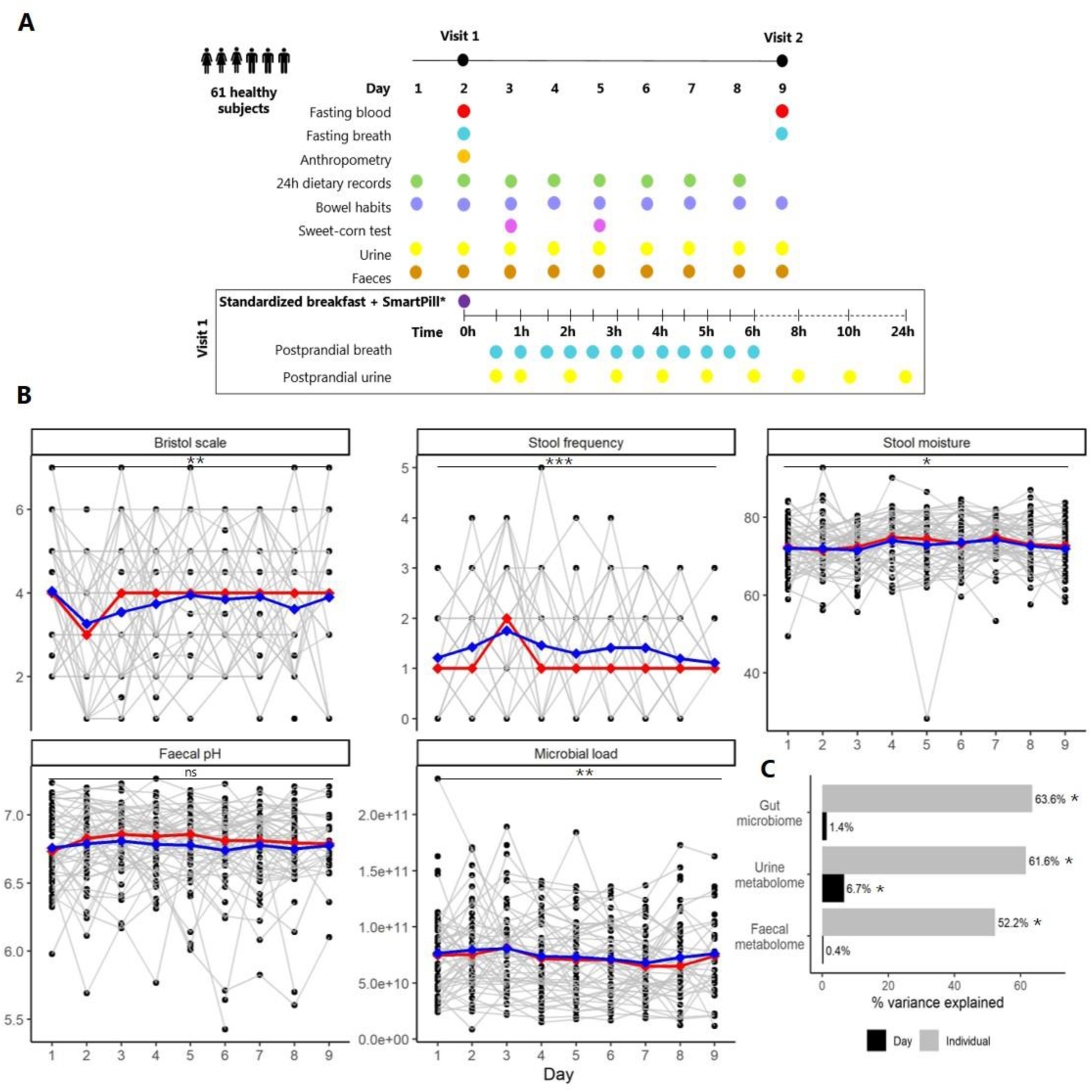
PRIMA study design and variations in gut environmental factors, gut microbiome and metabolomes. (A) PRIMA study design. The study included two site visits, at which fasting blood and breath samples were taken. At visit 1, anthropometric measurements were attained, and all participants were given a standardized breakfast and a subset of 50 volunteers ingested SmartPills immediately after. Postprandial breath hydrogen and methane were measured every 30 min for 6 h, and postprandial urine was collected at 0.5 h, and every hour until 24-h as indicated. On days 3 and 5, participants performed a sweet-corn test to measure whole gut transit time. In addition, daily 24-h dietary records (days 1 – 8), records of bowel habits (stool consistency, stool frequency, and time of defecation) as well as daily urine and faecal samples were obtained. (B) Inter- and intra-individual variations in the gut environmental factors over the 9 consecutive days. The red and blue lines represent median and mean values, respectively. Grey lines represent intra-individual fluctuations over time. Asterisks indicate the statistical significance of mixed-effect models accounting for repeated measures (***p-value < 0.001, **p-value < 0.01, *p-value < 0.05, ns; not significant, see **Supplementary Table 2** for details). (C) Percentage of variation explained by individual and study day in the gut microbiome, urine and faecal metabolome based on PERMANOVA tests (* p-value < 0.05).

**Table 1.** Participants’ characteristics (N = 61)

|  | Mean ± SD/<br>Median (25<sup>th</sup>-75<sup>th</sup> pct.) | Range |
| --- | --- | --- |
| Sex, male/female | 18/43 | - |
| Age, yr | 39 ± 13.5 | 20 - 66 |
| Body mass index, kg/m<sup>2</sup> | 23.6 ± 2.8 | 17.6 - 29.5 |
| Fasting glucose, mmol/L * | 5.1 (4.9 - 5.4) | 4.4 - 6.9 |
| Fasting insulin, mmol/L * | 32.5 (24.2 - 51.7) | 14.7 - 132.0 |
| Fasting C-pep, pmol/L * | 407 (321 - 520) | 186 - 771 |
| **Dietary intake\*** |  |  |
| Total energy intake, kcal/day | 2256 ± 605 | 1276 - 5091 |
| Carbohydrate, E% | 41.1 ± 8.7 | 14.2 - 69.4 |
| Protein, E% | 15.5 ± 4.4 | 5.5 - 35.6 |
| Fat, E% | 39.4 ± 8.2 | 16.5 - 62.3 |
| Fibre intake, g/1000kcal/day | 24.0 ± 10.3 | 3 - 62 |
| **Gut environmental factors\*** |  |  |
| Stool consistency, Bristol stool scale | 4 (3 - 5) | 1 - 7 |
| Stool frequency, n per day | 1 (1 - 2) | 0 - 5 |
| Stool moisture, % | 73 (69 - 77) | 28 - 93 |
| Faecal pH | 6.8 (6.3 - 7.0) | 5.4 - 7.3 |
| **Faecal SCFAs µmol/g of faeces\*** |  |  |
| Acetate | 16.11 (8.29 - 26.06) | 0.85 - 76.07 |
| Propionate | 2.49 (1.56 - 3.93) | 0.01 - 33.80 |
| Butyrate | 1.43 (0.78 - 2.09) | 0.04 - 5.96 |
| Valerate | 1.20 (0.89 - 1.69) | 0.33 - 4.10 |
| Caproate | 0.54 (0.13 - 1.07) | 0.01 - 6.88 |
| **Faecal BCFAs, µmol/g of faeces\*** |  |  |
| 2-methylbutyrate | 0.54 (0.39 - 0.71) | 0.07 - 3.24 |
| Isovalerate | 0.41 (0.28 - 0.55) | 0.08 - 2.23 |
| Isobutyrate | 0.29 (0.22 - 0.37) | 0.07 - 1.39 |
| **Breath\*** |  |  |
| Fasting hydrogen, ppm | 6.5 (4.0 - 12.0) | 0.5 - 51 |
| Fasting methane, ppm | 1.0 (0 - 18) | 0 - 67.5 |

\*-mean of all records/measurements, E%; energy percent, SCFAs; short-chain fatty acids, BCFAs; branched-chain fatty acids

The study included two visits (day 2 and day 9) at which fasting blood glucose, insulin and C-peptide as well as breath hydrogen and methane were measured (**Table 1**). On the first visit, all participants were given a standardized breakfast corresponding to 25 % of their daily energy demand (rye bread with butter and jam, boiled egg and yoghurt with blueberries and walnuts, **Supplementary Table 1**) before a subset of the volunteers (n = 50) ingested a wireless motility capsule (SmartPill®) to measure whole gut and segmental transit time and pH^15^. Subsequently, postprandial breath hydrogen and methane measurements (t = 30 min, 60 min, 90 min, 120 min, 150 min, 180 min, 210 min, 240 min, 270 min, 300 min, 330 min, and 360 min) and urine sampling (t = 30 min, 1 h, 2 h, 3 h, 4 h, 5 h, 6 h, 6-8 h, 8-10 h, 10-24 h) were obtained.

The participants recorded their food intake on a daily basis (day 1 – 8) using 24-h dietary records as implemented in the myfood24® nutrition platform (https://www.myfood24.org) with input from the National Danish Food database (https://www.frida.fooddata.dk). Furthermore, the participants recorded their daily bowel habits including stool consistency assessed by the Bristol stool scale (BSS)^16^, the time of defecation of each bowel movement, and stool frequency (number of bowel movements per day), and collected daily urine and faecal samples (the first bowel movement). The study population had normal bowel habits with BSS of type 4 (median, **Table 1**) and 1 bowel movement per day (median, **Table 1**).

In addition, transit time was also estimated by a self-administered sweet-corn transit time test^17^ on days 3 and 5. We measured faecal water content (indication of stool moisture, a proxy marker of transit time^17^), pH, and microbial load in all collected faecal samples (n = 484). In addition, all collected urine samples (daily spot samples and postprandial samples, n = 1154) and a subset of faecal samples (n = 170) were profiled by untargeted liquid chromatography-mass spectrometry (LC-MS)-metabolomics to obtain urine and faecal metabolomes. Finally, we obtained the gut microbiome composition via 16S rRNA gene sequencing of a subset of faecal samples (n = 362) and assessed both relative microbiome profiles (RMP) and quantitative microbiome profiles (QMP) after adjustment for the microbial load as previously described^18^.

### Variations of gut environmental factors, gut microbiome and metabolomes over time

Daily sampling allowed us to evaluate the fluctuations of the gut environmental factors, faecal- and urine metabolomes, gut microbiomes and diets within and between healthy adults over time **(Supplementary Fig. 2).**

Firstly, we observed that the gut environmental factors vary in the extent of their day-to-day fluctuations within individuals (**Figure 1B)**. The coefficient of variation within individuals (**Supplementary Table 3**) was ranging 0.3 –8.1 % for faecal pH, 0 – 57.8 % for BSS, 0 - 73.1 % for stool frequency, 2.2 – 24 % for stool moisture, and 7.6 –72.7 % for microbial load, suggesting that some individuals are more stable in terms of their gut environment than others. Moreover, on average all of the gut environmental factors including BSS, stool frequency, stool moisture, and microbial load fluctuated over the 9 days, whereas faecal pH remained stable **(Figure 1B, Supplementary Table 2A)**. In addition, Participant ID significantly accounted for day-to-day fluctuations in all of the gut environmental factors (**Supplementary Table 2B**), indicating that the gut environment is to some extend personal.

Next, we performed a permutational multivariate analysis of variance (PERMANOVA) on the gut microbiome (QMP), urine metabolome, and faecal metabolomes and found that the individual explained more than 50 % of the inter-individual variations in all three cases (**Figure 1C**). In contrast, the sampling day explained on average 6.7% of the urine metabolome variation but did not explain day-to-day variations in the gut microbiome and faecal metabolome (**Figure 1C**). Nonetheless, by inspecting the β-diversities of individual microbiome and metabolome profiles, we could see that some individuals showed less variation over the study period than others (**Supplementary Fig. 3**).

### Stool moisture and faecal pH contribute to intra-individual fluctuations in the gut microbiomes and metabolomes

To explore what drives the intra-individual fluctuation in the metabolomes and the microbiome, we performed distance-based redundancy analysis (db-RDA) with dietary macronutrients (carbohydrates, proteins, fats), fibres, coffee, and alcohol, all previously linked to the gut microbiome^1,3,19^ as well as the gut environmental factors (BSS, stool frequency, time of defecation, faecal pH, stool moisture). While none of the dietary components significantly explained intra-individual fluctuations in the gut microbiome or metabolomes, stool moisture, faecal pH, BSS, and time of defecation had significant effects on the gut microbiome (QMP, genus level, **Figure 2A**). Stool moisture, BSS, and time of defecation explained 3.5 %, 2 %, and 1.3 %, respectively, in line with previous reports^6,20^. All of these factors are proxies for gut transit time, suggesting that day-to-day variations in transit time are reflected in the gut microbiome variation. Additionally, faecal pH explained 2.5 % of the QMP data variation. In comparison, stool moisture, and faecal pH were also significant contributors to variation when analyzing RMP, but BSS and time of defecation were not (**Supplementary Fig. 4A**).

**Figure 2.**
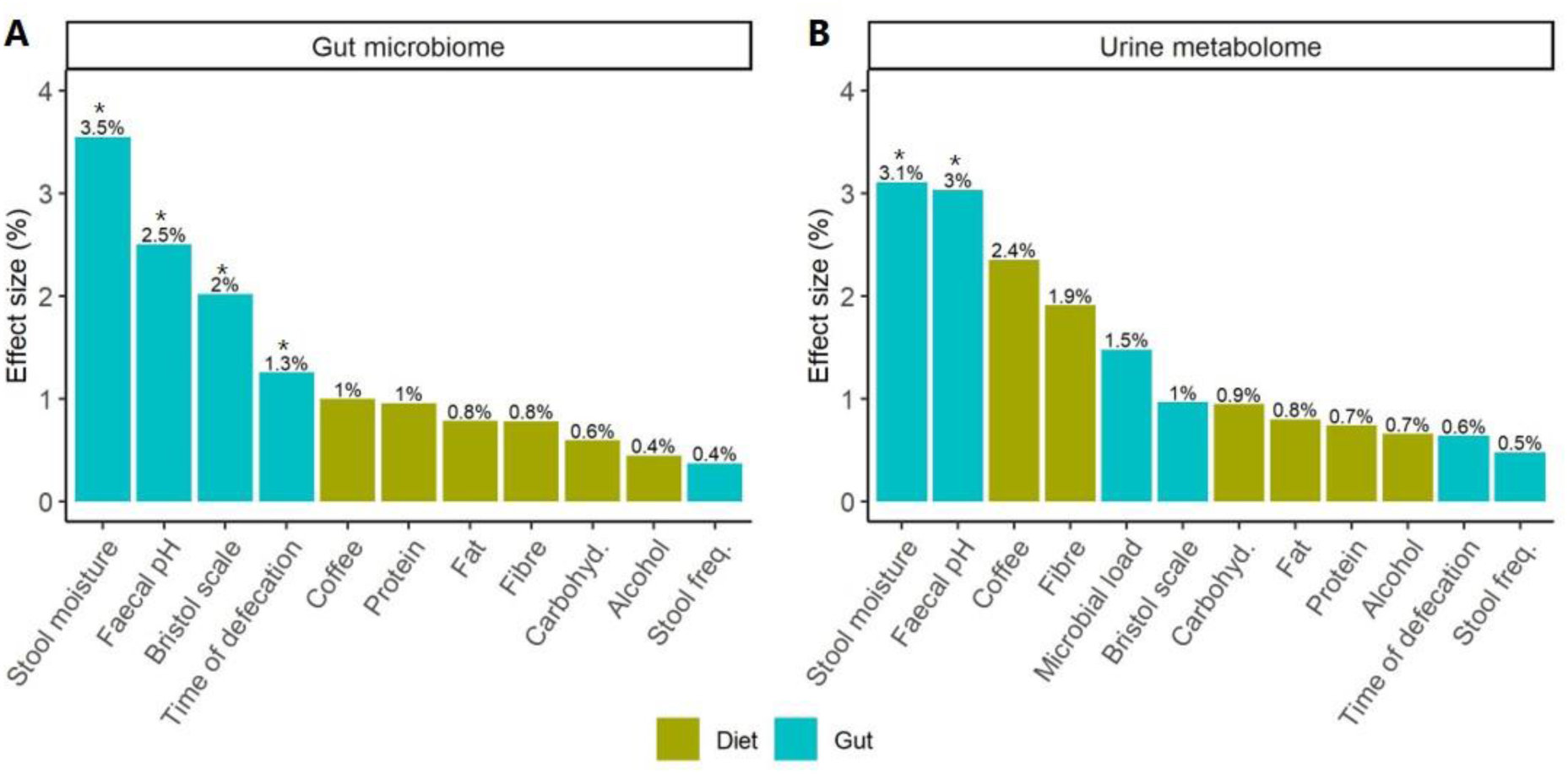
Contributions of dietary and gut environmental factors on intra-individual variations in (A) gut microbiome and (B) urine metabolome. The analysis was performed with distance-based redundancy analysis (db-RDA) with permutation test on daily quantitative microbiome data (QMP) and untargeted urine metabolome data with Bray-Curtis distances. The asterisks indicate statistical significance (*q-value < 0.05). See **Supplementary Fig. 4A and 4B** for relative microbiome profiles and faecal metabolome.

Stool moisture and faecal pH further significantly explained 3.1 % and 3 %, respectively, of the intra-individual variation in the urine metabolomes, despite both variables being quite stable over time (**Figure 2B**). This implies that even subtle changes in the colonic water content and pH may impact the host-microbiota metabolism reflected in the urine. Nevertheless, gut environmental factors did not significantly contribute to the intra-individual fluctuations in the faecal metabolomes (**Supplementary Figure 3B**). In this context it should be noted that faecal metabolome data were derived only from three consecutive days (spanning the SmartPill ingestion) and stool moisture still tended (p = 0.081) to have an effect.

### Inter-individual variations in the whole gut and segmental transit times and pH

The use of ingestible SmartPills allowed us to obtain whole gut transit time (WGTT) and segmental transit times including gastric emptying time (GET), small bowel transit time (SBT), colonic transit time (CTT), and intestinal transit time (ITT, SBT + CTT) as well as pH throughout the GIT (**Methods, Supplementary Fig. 4C**). We noticed in 8 individuals that the capsule was retained in the stomach for more than 8 hours, which is a common event also reported in other studies^21,22^. Therefore, GET and WGTT values from participants with GET > 8 h were excluded from the statistical analyses (n = 8). Furthermore, CTT and WGTT could not be determined in one participant due to the loss of signal to the receiver during the passage.

The median values of the segmental transit times were as follows; GET 4.8 h (range 3.1 - 6.2 h), WGTT 23.3 h (12.4 - 72.3 h), CTT 13.6 h (2.1 - 63.5 h), and SBT 5.1 h (2.5 - 10.3 h), which are in agreement with previously reported data on healthy populations^23^. For comparison, we also included two self-administered sweet-corn transit time assessments (i.e. corn TT, on days 3 and 5) with a median of 23.56 h (10.8 - 109.7 h) at day 3 and 19.7 h (12.0 - 84.5 h) at day 5. The median of the mean corn transit time across the two days was 21.72 h (11.75 – 97.08 h) (**Figure 3A**), similar to the WGTT obtained by the SmartPill. Additionally, the median coefficient of intra-individual variation for the corn TT was 18.2% (range 0.4 – 77.9%) and we found a strong correlation between the two measurements (Spearman correlation coefficient (SCC) = 0.8, p < 0.001) suggesting consistency between the days within individuals. Notably, we did not observe any correlation between WGTT or CTT and the corn TT (**Supplementary Fig. 5**) indicating that despite providing similar results on average, individually, these two methods showed different results. Yet, when exploring the relationships between segmental transit times, proxy markers, gut environmental factors and subject characteristics (**Supplementary Fig. 5),** we found that the transit times recorded by both methods were negatively correlated to BSS and stool moisture, in agreement with previous reports^17,24^.

**Figure 3.**
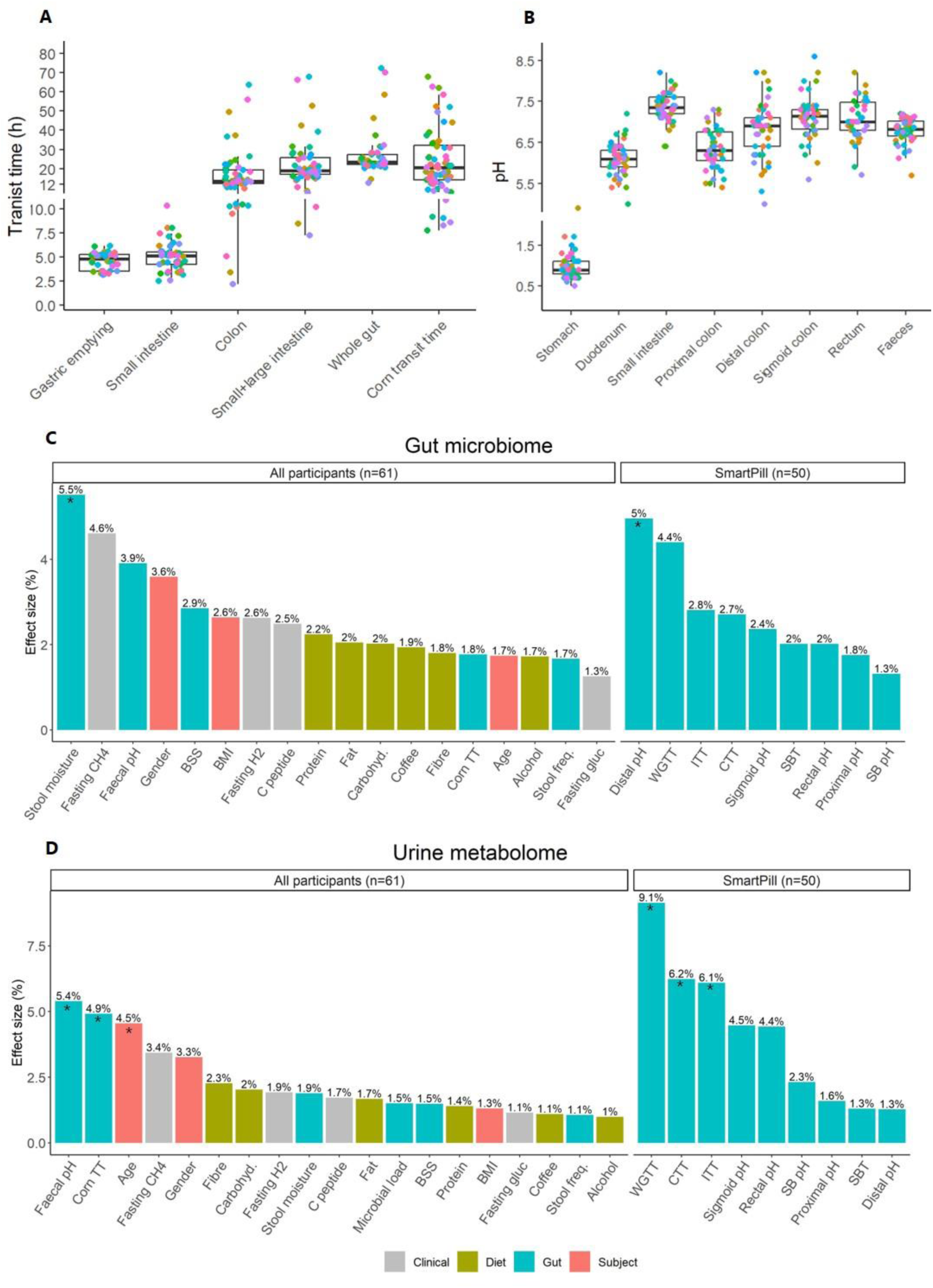
Variations in the whole gut and segmental transit times and pH and their effect on the inter-individual variability in the gut microbiome and urine metabolome. (A) Boxplots showing segmental and whole gut transit time measured by the SmartPill (n = 50) at day 2 and mean transit time of sweet-corn (n = 61, day 3 and 5) with each dot representing an individual. (B) Boxplots showing pH throughout the gastrointestinal tract measured by the SmartPill and in faeces measured by pH meter at day 2 with each dot representing an individual. (C) Contributions of clinical variables, dietary components, gut environmental factors, and subject characteristics to inter-individual variations in the gut microbiome (QMP, sample closest to the capsule body exit) urine metabolome (24-h, day 2), respectively, quantified by distance-based redundancy analysis with permutation tests. Effect sizes are plotted. The asterisks indicate statistical significance (*q < 0.1). SB; small bowel; SBT; small bowel transit time, CTT; colonic transit time, WGTT; whole gut transit time, BMI; body mass index, gluc; glucose, BSS; Bristol stool scale; CH4; methane, H2; hydrogen.

Large inter-individual variations in the gastrointestinal segmental pH were observed (**Figure 3B**) with the following pH values in the upper GIT; the stomach (median 0.9, range 0.5 - 4.9), duodenum (6.1, 5.0 - 7.2), and small intestine (7.4, 6.4 - 8.2). pH in the proximal colon was slightly acidic (6.3, 5.3 - 7.0) followed by a gradual pH increase in the distal colon 6.9 (5.0 - 8.2) and sigmoid colon 7.2 (5.6 - 8.6). Interestingly, a small decrease in pH was observed from the sigmoid colon to the rectum (7.0, 5.7 - 9.6) and also in the faecal pH (6.9, 6.6 - 7.3) indicating that acidifying processes occur after entry into the rectum.

### Colonic transit time and pH contribute to inter-individual variations in the gut microbiome and metabolomes

To quantify the degree by which subject characteristics (i.e. age, sex, and BMI), clinical variables (fasting glucose and C-peptide, breath measurements), diet (mean intake of carbohydrates, proteins, fats, fibres, coffee and alcohol), and gut environmental factors explain inter-individual variations in the gut microbiome and the metabolomes, we performed db-RDA with permutation tests on data derived from faecal samples and 24-h urine collected on day 2 for all participants (n = 61, **Supplementary Table 4**). Moreover, we performed the same analysis with whole gut and segmental transit times and pH derived from the SmartPills on day 2 (n = 50).

As seen for the intra-individual QMP fluctuations, we found that stool moisture and distal colon pH were important factors associated with the inter-individual variation in QMPs (**Figure 3C**) accounting for 5.5 % and 5 % of the variation, respectively, on day 2. Importantly, stool moisture and distal pH also showed significant contributions to the inter-individual variation in QMP on other days as well (**Supplementary Table 4**). But unlike previously reported data from larger cohorts, BSS did not explain a significant proportion of the microbiome variation in this population.

WGTT, CTT, corn TT, and faecal pH explained 9.1 %, 6.2%, 4.9 %, and 5.4 %, respectively, of the inter-individual variations in the 24-h urine metabolome, in addition to age, which explained 4.5 % of the variation (**Figure 3D**). These contributions were consistent when testing against the urine metabolomes on different days (**Supplementary Table 4**). On the contrary, segmental transit time did not significantly contribute to the inter-individual variation in the faecal metabolomes, whereas pH in the distal colon, alcohol and fibre intake showed the largest effects explaining 6.8 %, 6.2 %, and 5.9 % respectively, however this was not significant after adjusting for multiple testing (**Supplementary Table 4**).

Altogether, these results emphasize that the personal environment in the gut contributes considerably to the inter-individual differences not only in the gut microbiota but also in the urinary metabolic profiles.

### Intra-individual fluctuations in microbial-derived metabolites and their associations to gut environmental factors and diet

We assessed intra-individual fluctuations in breath hydrogen and methane between days 2 and 9 (**Table 1**, **Figure 4A, Supplementary Fig. 6A**). Additionally, concentrations of SCFAs, namely acetate, propionate, butyrate, valerate, and caproate, and BCFA, namely isobutyrate, isovalerate, and 2-methylbutyrate, were quantified by LC-MS targeted analysis in all faecal samples collected between the SmartPill ingestion and egestion for each subject (n = 170) (**Table 1**, **Figure 4B and 4C, Supplementary Fig. 6B and 6C**). In agreement with previous data, acetate was found in the highest concentrations (median 16.11 µmol/g of faeces, range 0.85 - 76.07), followed by propionate (2.49 µmol/g of faeces, 0.01 - 33.80) and butyrate (1.43 µmol/g of faeces, 0.04 - 5.96) in all subjects. For the majority of the subjects, 2-methylbutyrate (0.54 µmol/g of faeces, 0.07 - 3.24) was the most abundant BCFA.

**Figure 4.**
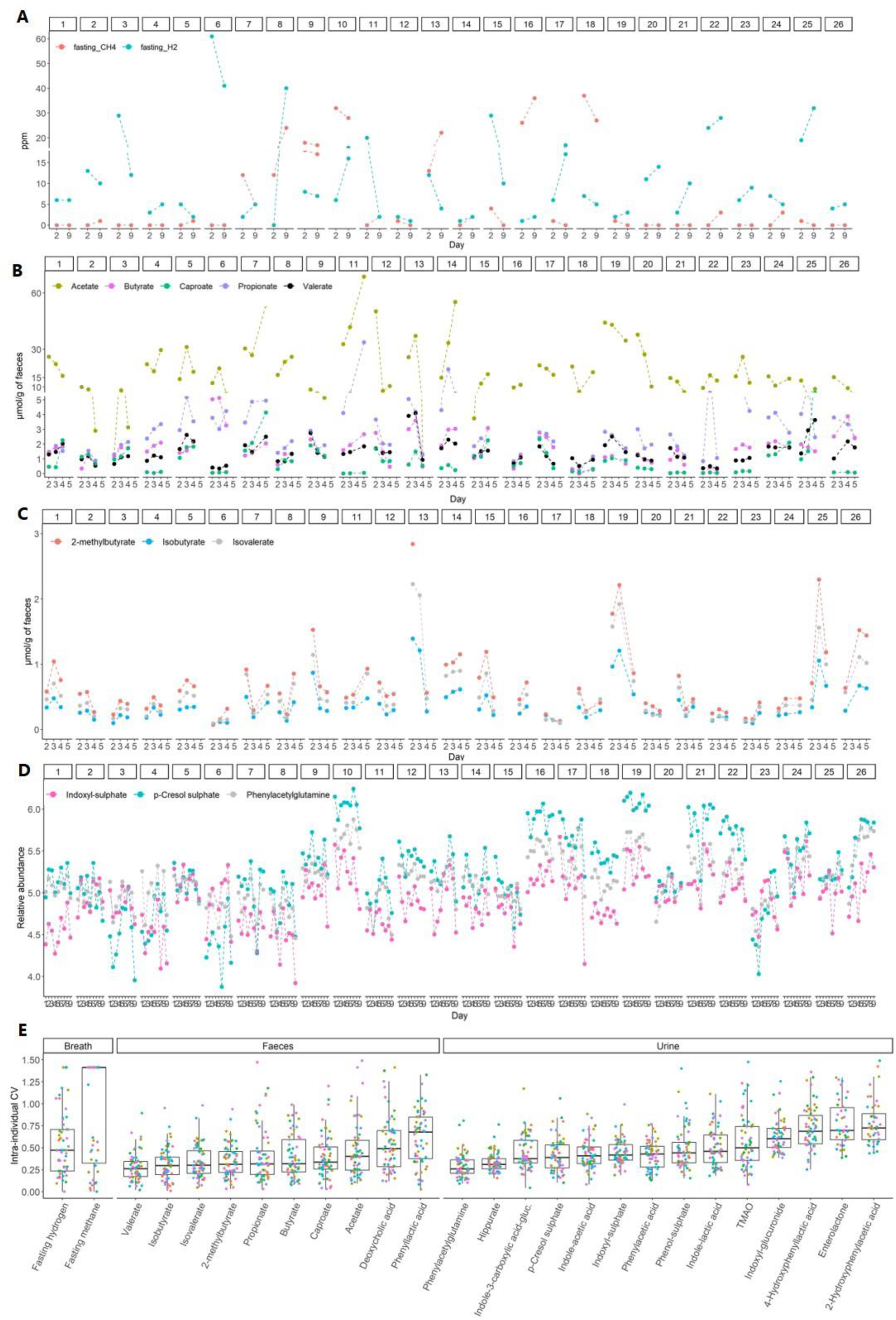
Concentrations and intra-individual variations in microbial-derived metabolites measured in breath, faeces and urine. (A) Fasting breath levels of hydrogen and methane (ppm) on days 2 and 9 for the first 26 individuals. (B) Faecal concentrations of short-chain fatty acids (SCFAs) for 26 selected individuals over 4 days. (C) Faecal concentrations of branched chain-fatty acids (BCFAs) for 26 selected individuals over 4 days. (D) Relative abundances of urinary markers of microbial proteolysis for 26 selected individuals over 9 days. See **Figure S5** for profiles of all 61 study participants. (E) Boxplots showing coefficients of intra-individual variations for microbial-derived metabolites measured in breath, faeces and urine. Each dot represents an individual.

**Figure 5.**
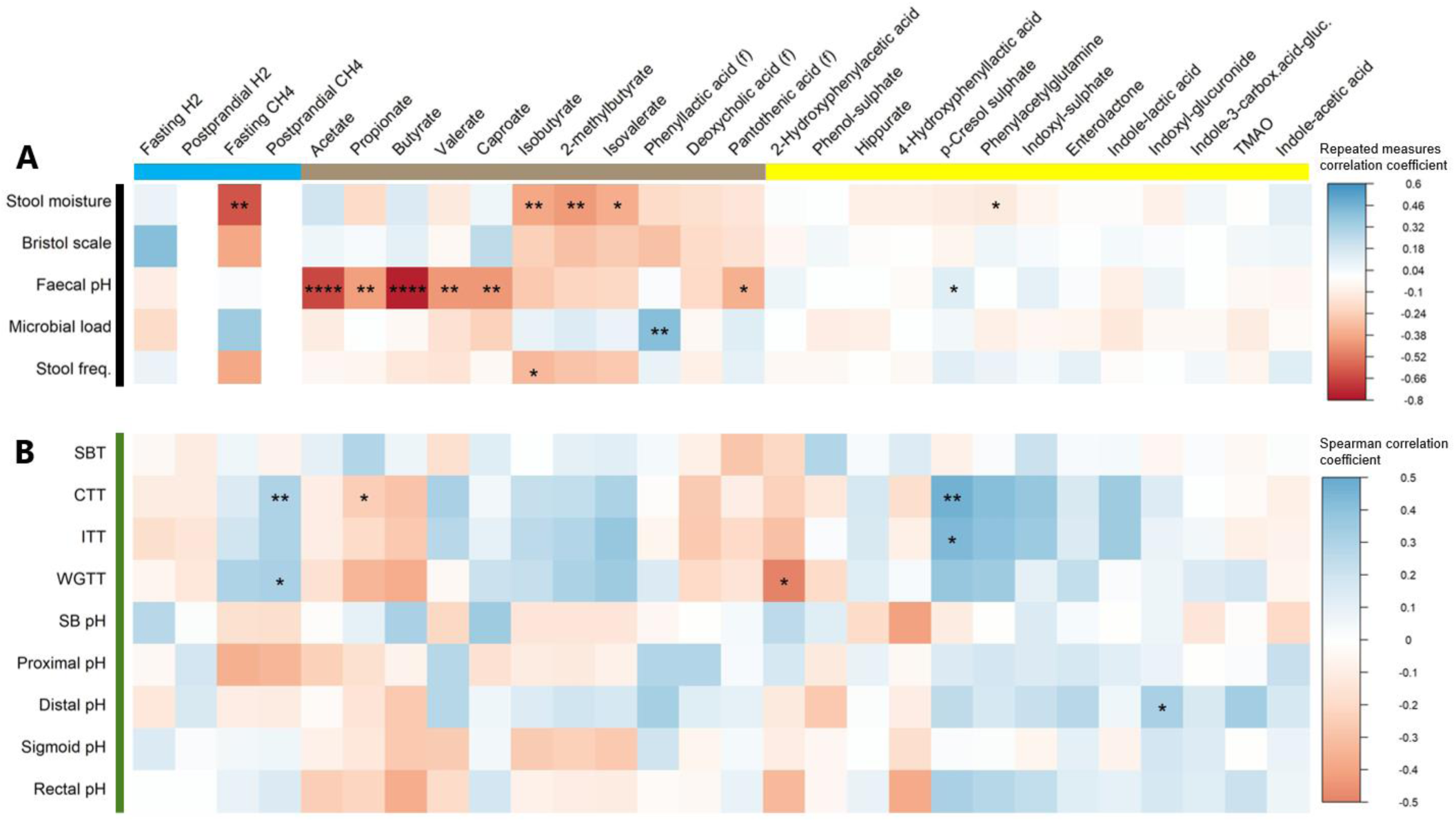
Correlation analysis between gut environmental factors, segmental transit times and pH assessed by the SmartPill and microbial-derived metabolites. **(A-B)** The colour gradient shows (A) repeated measures correlation coefficient or (B) the Spearman correlation coefficient and the asterisks indicate statistical significance (****q < 0.001,*** q < 0.01,** q < 0.05,*q < 0.1). Blue, brown, and yellow bars indicate breath, faecal, and urine metabolites, respectively. The black bar indicates repeated measure correlations where daily values for each variable have been used (panel A, **Supplementary Fig. 7**), whereas the green bar indicates analysis based on data collected on day 2 (panel B). Postprandial hydrogen and methane were only measured at one-time point and therefore were not included in the repeated measure analysis. CH4; methane, CTT; colonic transit time, (f); faecal, ITT; intestinal transit time, H2; hydrogen, SB; small bowel, SBT; small bowel transit time, TMAO; trimethylamine N-oxide; WGTT; whole gut transit time

Secondly, microbial-derived metabolites represented in our in-house collection of reference compounds were identified in the obtained faecal and urine metabolomes. We were particularly interested in proteolytic markers, including p-cresol sulphate, phenylacetylglutamine, and indoxyl sulphate, since we have previously linked these to inter-individual variations in gut transit time^9,17,25^. All three proteolytic markers were detected in all urine samples from the 61 subjects (**Figure 6D**, **Supplementary Fig. 6D**). In addition, we identified seventeen other microbial-derived metabolites in the urine. We calculated the CV for intra-individual variations (CV_Intra_) of the identified metabolites in breath, faeces and urine among all participants. Substantial differences in intra-individual variations were observed (**Figure 4E**). Breath methane and hydrogen had a median CV_Intra_ of 141 % and 47 %, respectively, yet we found a moderate positive correlation between the two-time points for both gases (hydrogen: SCC = 0.42, p < 0.001; methane: SCC = 0.66, p < 0.001). Moreover, faecal concentrations of the SCFAs and BCFAs fluctuated considerably from day-to-day (median CV_Intra_ ranging from 26 % to 40 %) with valerate varying the least and acetate the most. Similarly, the relative abundances of the measured metabolites in urine varied substantially from day to day with a median CV_Intra_ of 26 %, 42 % and 39 % for phenylacetylglutamine, indoxyl sulphate, and p-cresol sulphate, respectively. These findings suggest that microbial-derived metabolites in breath, faeces and urine fluctuate from day-to-day on a habitual diet.

**Figure 6.**
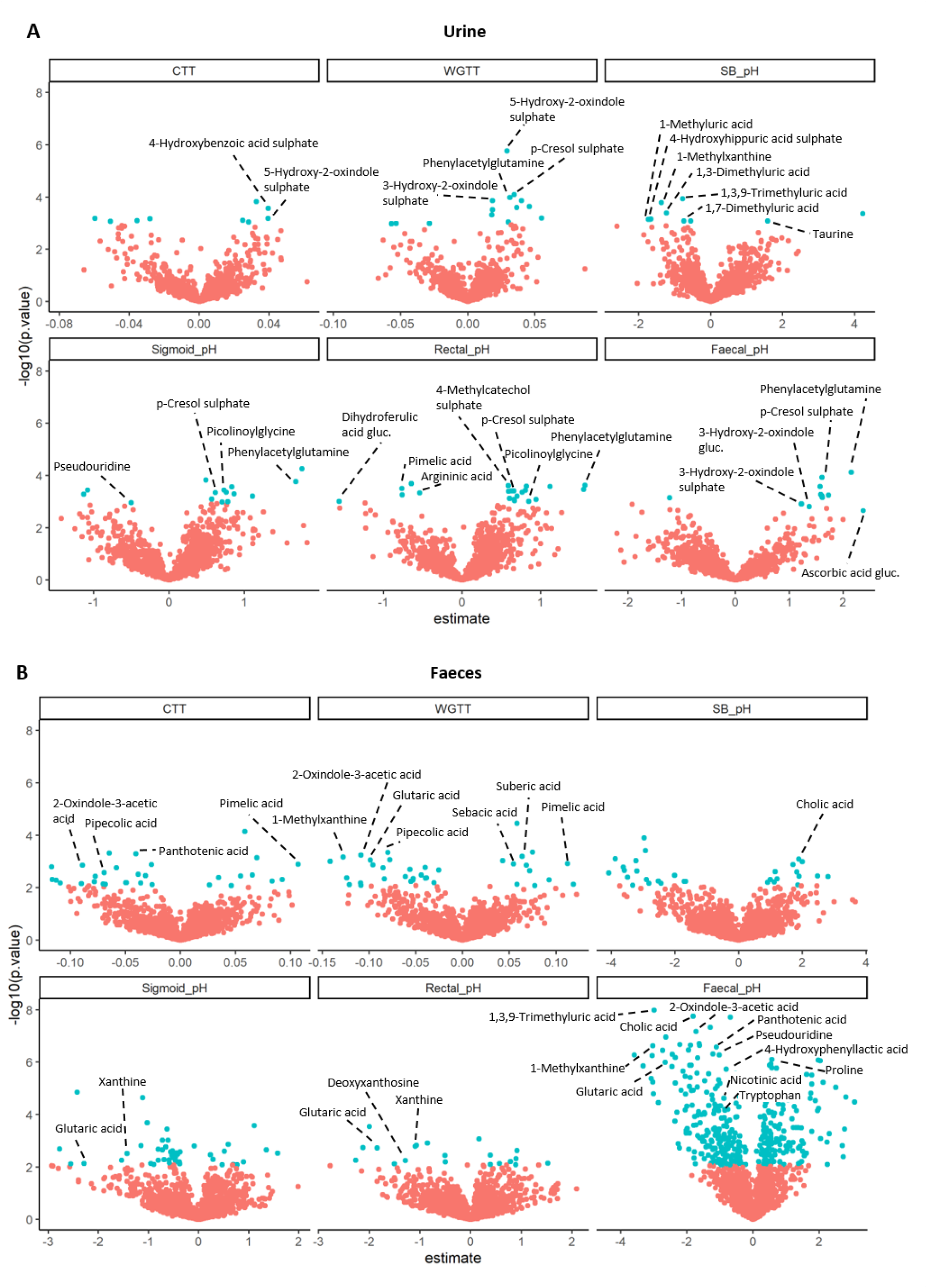
Metabolites identified via LC-MS untargeted metabolomics associated with segmental transit time and pH. **(A-B)** Volcano plots derived from regression models where each dot represents a metabolic feature with blue representing statistically significant associations (q < 0.1) in (**A**) urine and (**B**) faeces. The x-axis shows the regression coefficient values (estimate) indicating either positive or negative associations. CTT; colonic transit time, SB; small bowel, WGTT; whole gut transit time

Given that gut environmental factors explained large proportions of the gut microbiome and urine metabolome intra- and inter-individual variations, we hypothesized that specific metabolites would be linked to these factors. Indeed, by correlating gut environmental factors with microbial-derived metabolites using repeated measurements (**Figure 5A and Supplementary Fig. 7**), we found several significant associations. In particular, stool moisture was negatively correlated to several markers of microbial proteolysis including urinary phenylacetylglutamine (r = −0.12, q < 0.1), faecal isobutyrate (r = −0.39, q < 0.05), isovalerate (r = −0.37, q < 0.1), and faecal 2-methylbutyrate (r = −0.43, q < 0.05) over days, similar to a previous cross-sectional study^26^. Fasting breath methane was negatively associated with stool moisture (r = −0.62, q < 0.05) on both days, indicating increased methanogenesis with longer colonic transit time reflected by lower stool moisture. Moreover, daily faecal pH was positively correlated to urinary p-cresol sulphate (r = 0.12, q < 0.1) in contrast to faecal SCFAs that showed a negative correlation to faecal pH with butyrate showing the strongest correlation (r = −0.77, q < 0.001) in line with previous human studies^27^. In addition, phenyllactic acid was positively correlated to microbial load (r = 0.42, q < 0.05). Taken together, our findings suggest that slower colonic transit and/or higher pH are linked to increased microbial proteolysis and methanogenesis as opposed to microbial saccharolysis resulting in SCFAs that are consistently associated with lower colonic pH.

Since diet provides the substrates for gut microbial metabolism, we performed the same analysis exploring the relationships between daily intake of macronutrients, dietary fibres, and microbial-derived metabolites (**Supplementary Table 5**). We found positive associations between the urinary levels of hippurate and the intake of proteins (r = 0.17, q < 0.05) but not fruits and vegetables^28,29^. Interestingly, we also observed a tendency for a negative association between intake of starch and urinary levels of TMAO (r = −0.13, p = 0.02, q = 0.09), and between intake of dietary fibres and several proteolytic metabolites including all three faecal BCFAs (isobutyrate: r = −0.32, p = 0.02, q = 0.14, isovalerate: r = −0.31, p = 0.03, q = 0.14, 2-methylbutyrate: r = −0.29, p = 0.04, q = 0.18), urinary p-cresol sulphate (r = −0.11, p = 0.04, q = 0.09), and phenylacetylglutamine (r = −0.12, p = 0.03, q = 0.09).

### Inter-individual variations in microbial-derived metabolites and their associations to segmental transit time and pH

Several microbial-derived metabolites were associated with gut environmental factors measured in faeces longitudinally over 9 days. Therefore, we next wanted to explore links between the microbial-derived metabolites and whole gut and segmental transit times and pH (**Figure 5B**) measured on day 2. Spearman correlation analysis showed that a shorter CTT was significantly associated with higher faecal propionate levels (SCC = −0.25, q < 0.1) and a similar trend was observed with faecal butyrate (SCC = −0.29, p < 0.05). Notably, the same trends for SCFAs were observed with ITT and WGTT but not with SBT. A tendency for a negative correlation between faecal butyrate and rectal pH was also observed (SCC = −0.37, p < 0.05), but not with pH in other segments of the colon, suggesting that butyrate production may contribute to the reduced pH observed in rectum and in faeces. The correlation was the opposite between faecal butyrate and pH in the small intestine (SCC = 0.33, p < 0.05) and no correlations were found between other faecal SCFAs and small intestinal pH.

Furthermore, longer CTT and ITT were associated with significantly higher urinary levels of p-cresol sulphate (SCC = 0.48, SCC = 0.44, respectively, q < 0.05) and similar tendencies were found with phenylacetylglutamine (SCC = 0.43, SCC = 0.40, p < 0.01), indoxyl-sulphate (SCC = 0.39, SCC = 0.36, p < 0.05) and indole-lactic acid (SCC = 0.36, SCC = 0.35, p < 0.05), which was not observed for SBT indicating that CTT determines the abundance of the proteolytic metabolites. These findings support the hypothesis that longer passage through the colon is linked to microbial proteolysis possibly due to the depletion of substrates for saccharolytic fermentation^9,30^. In line herewith, indoxyl-glucuronide was positively associated with pH in the distal colon (SCC = 0.33, q < 0.1) and a similar trend between urinary p-cresol sulphate and rectal pH (SCC = 0.32, p < 0.05) was observed. The same tendencies were found for other proteolytic markers emphasizing that microbial proteolysis is linked to higher colonic pH. Notably, the correlations between proteolytic markers and pH were stronger with pH in the distal colon than with pH in the small intestine and the proximal colon indicating higher contribution of microbial proteolysis to pH in the distal gut compared to the proximal gut.

Positive correlations were found between postprandial methane and CTT (SCC = 0.37, q < 0.05), ITT (SCC = 0.4, p < 0.05), and WGTT (SCC = 0.32, p < 0.05), but not SBT with the same tendencies for fasting breath methane. In summary, these results show that CTT and colonic pH but not SBT and small intestinal pH are associated with levels of microbial-derived metabolites in breath, faeces and urine.

### Untargeted metabolomics revealed novel associations with segmental transit time and pH

Untargeted metabolomics is a powerful tool that can be applied to study host-microbiota interactions. To explore unknown metabolic features related to the gut environment, we employed univariate and multivariate statistical models on all molecular features identified in the urine (n = 641 in positive mode, n = 651 in negative mode) and faeces (n = 453 in positive mode, n = 445 in negative mode). Firstly, we used sparse partial least squares (SPLS) models using the SmartPill-derived data and urine metabolomes from 24-h postprandial urine that was collected on day 2. Similarly, faecal metabolomes collected closest to the SmartPill egestion were used. Secondly, we performed linear regression models on the same data and further investigated features selected by both models (446 unique features).

Apart from urinary levels of p-cresol sulphate and phenylacetylglutamine positively associated with WGTT, sigmoid, rectal and faecal pH, several metabolic features in urine and faeces were associated with whole gut and segmental transit times and pH (**Figure 6A and 6B)**. To investigate these features in further detail, the corresponding samples were analyzed by tandem MS and by matching with authentic standards when available, resulting in the identification of 33 metabolites **(Supplementary Table 6 and 7)**. However, a large number of features (n = 382) remain unidentified.

In urine, we identified 5-hydroxy-2-oxindole sulphate, 3-hydroxy-2-oxindole sulphate, and 4-hydroxybenzoic acid sulphate to be positively associated with WGTT and/or CTT. Moreover, 3-hydroxy-2-oxindole glucuronide was positively correlated to faecal pH. Similar to p-cresol sulphate and phenylacetylglutamine, these metabolites likely originate from microbial catabolism of the aromatic amino acids tryptophan and tyrosine, emphasising the link between longer transit time/higher faecal pH and increased microbial proteolysis. In support of this, faecal tryptophan levels were negatively linked to faecal pH potentially indicating that tryptophan is being less utilized by the gut microbiota with shorter transit time and/or when carbohydrates are available. Moreover, higher levels of amino acid proline in faeces and picolinoylglycine in urine were associated with increased faecal and rectal pH, respectively.

Several dicarboxylic acids in faeces, pimelic (C7) suberic (C8), and sebacic acid (C10) were positively associated with WGTT and CTT. Pimelic acid and suberic acid may originate from microbial metabolism of fatty acids (e.g. oleic acid)^31,32^, which may imply that the excretion of microbial metabolites derived from dietary fats increases with increasing transit time. Faecal glutaric acid (C4) was, however, negatively correlated with WGTT, sigmoid, rectal and faecal pH. We also identified pipecolic acid in faeces, which was negatively associated with WGTT and CTT. Pipecolic acid is highly abundant in plants, however, it can also be produced by the gut microbiota from lysine^33^. Furthermore, higher urinary levels of citric acid were positively associated with pH in the proximal colon.

Moreover, faecal levels of 2-oxindole-3-acetic acid, previously linked to the New Nordic diet and Mediterranean diet^34,35^, were negatively associated with WGTT, CTT, and faecal pH. Similarly, faecal pantothenic acid and vitamin B, nicotinic acid (B3, niacin), were negatively associated with CTT and/or faecal pH. Additionally, there was a negative correlation between dihydroferulic acid glucuronide and argininic acid in urine and rectal pH, while p-hydroxyphenyllactic acid in faeces was negatively linked with fecal pH.

4-Hydroxyhippuric acid and several urinary markers of coffee intake were negatively associated with pH in the small intestine, namely 1-methyluric acid, 1-methylxanthine, 1,3-dimethyluric acid, 1,7-dimethyluric acid, and 1,3,9-trimethyluric acid. Faecal 1-methylxanthine and 1,3,9-trimethyluric acid were also negatively associated with WGTT and/or faecal pH indicating that coffee consumption might be linked to both, intestinal pH and transit time. We also found a positive association between rectal pH and urinary 4-methylcatechol sulphate, a microbial metabolite of quercetin, found in many plant-based foods^36^. In addition, taurine in urine and a cholic acid in faeces were positively associated with small intestinal pH, suggesting a link between bile acids and small intestinal pH in agreement with the fact that secretion of bile into the small intestine neutralizes the acidic chyme from the stomach^37^.

Finally, in line with our previous work where the urinary level of pseudouridine was inversely associated with CTT^9^, we found a negative association between pH in the sigmoid colon and urinary pseudouridine, a primary constituent of RNA. Similarly, pseudouridine was also identified in faeces where it showed an inverse relationship to faecal pH as did deoxyxanthosine and xanthine, suggesting that increased cell turnover is linked to lower colonic pH.

Altogether, by employing untargeted LC-MS metabolomics, we identified several RNA-, microbial-, and food-derived metabolites newly associated with WGTT, CTT and pH in the distal part of the colon emphasizing an interplay between the gut environment and the diet-microbiota interactions.

## Discussion

Gut transit time and pH are important determinants of gut microbiota composition and metabolism^7^. Here, we showed that the gut environment also significantly explains both intra- and inter-individual variations in the gut microbiome composition and host-microbiota co-metabolism, as reflected by associations to microbial-derived metabolites measured in breath, faeces and urine.

In this study, we demonstrated that whole gut and segmental transit time and pH measured by the SmartPills as well as the sweet-corn test substantially varied between healthy individuals. The considerable differences in luminal pH among individuals can introduce challenges in studies using ingestible sampling devices that rely on pH sensitivity to target specific regions of the gut^38,39^. Furthermore, these insights are of fundamental importance since pH and transit time are key factors for shaping microbial growth and enzyme activities^40^. Therefore, the regional variations in pH and transit time could potentially be key for shaping the regional microbiome and metabolism along the GIT, and potentially explain inter-individual differences in the gut microbiome composition and microbiome-responses to foods. In support hereof, distal colonic pH and WGTT contributed the most to faecal microbiome and metabolome variation. Future studies sampling along the GIT combined with measurements of regional pH and transit time are needed to ultimately disentangle this. Recently, a study using ingestible sampling devices^38^ showed indeed that microbiome and metabolome compositions differ along the GIT.

The daily sampling during the 9 days allowed us to follow day-to-day fluctuations in microbial-derived metabolites. All of the microbial-derived metabolites in this study showed large variations both between and also within individuals, which highlights the importance of the need for repeated measurements in human study designs. We showed that the variations in many microbial-derived metabolites were linked to gut environmental factors with a strong link between longer transit time and increased levels of metabolites derived from microbial proteolysis including p-cresol sulphate. Interestingly, more than 40 % of plasma variation in p-cresol sulphate has previously been explained by microbiome variation^41^, suggesting that the inter-individual differences in this metabolic feature might largely be confounded by differences in bowel habits. In fact, many of the metabolites associated with longer transit time in our study have been reported to be elevated in patient groups with constipation^42–44^ demonstrating the need for transit time estimates in future microbiome studies. Based on our data, stool frequency was an insensitive measure for a small cohort with healthy individuals whereas stool moisture appeared as a significantly more informative proxy marker.

By untargeted metabolomics, we discovered aromatic amino acid-derivatives and dicarboxylic acids including pimelic acid, which have not previously been linked with intestinal transit time and/or pH. Pimelic acid can originate from microbial metabolism of fatty acids^31,32^ and has previously been found at elevated faecal levels in patients with chronic kidney disease^45^ and colorectal cancer^46^; who often suffer from constipation^47,48^. Moreover, pimelic acid can be synthesized by the gut microbiota as a part of biotin (vitamin B7) synthesis^31^ and biotin deficiency has been linked to obesity^49^ and irritable bowel disease^49^. Recently, metabolic profiling of samples collected along the gastrointestinal tract in humans has shown that the abundance of dicarboxylic acids increases towards the distal gut^38^. The authors hypothesized that this might be due to the catabolism of host epithelial cells, which combined with our results suggests that longer intestinal transit time might be associated with increased shedding of epithelial cells into the intestine. However, additional research is needed to better understand the interplay between dicarboxylic acids, gut environment and host physiology.

Besides dicarboxylic acids, we found that faecal nicotinic and pantothenic acids showed an inverse relationship to faecal pH. Pantothenic acid, also known as vitamin B5, is synthesized by the gut microbiota and has previously been linked to dietary fibre intake^50^ and significantly decreased faecal levels were found in patients with Parkinson’s disease^19^. Nicotinic acid can be synthesized by gut microbiota from tryptophan and contributes to gut homeostasis^51^. As higher faecal pH is positively correlated to a longer transit time, a longer transit time may imply lower tryptophan availability in the gut for the synthesis of nicotinic acid. Nicotinic acid, as well as pantothenic acid, were both previously found at decreased faecal levels in patients with ulcerative colitis when compared with healthy individuals^52^. However, since dietary wholegrains and fibres are rich in both vitamins^53^ the inverse relationship to faecal pH might originate from decreased carbohydrate availability in the colon with a longer passage. In addition, we identified several coffee-derived metabolites associated with lower small intestinal pH including 1-methylxanthine, which was also linked to shorter WGTT. Previous studies have shown that coffee stimulates colonic motility possibly due to the release of gut hormones regulating motility and/or localized effects of specific coffee-derived metabolites^54–56^, yet this needs further elucidation.

Collectively, our data support previous findings suggesting a shift in microbial metabolism from saccharolysis (carbohydrate fermentation) towards utilization of other substrates (i.e. proteins and lipids) with longer passage through the colon^9,30^.

Curiously, we found a negative association between intake of daily dietary fibres and several of the proteolytic markers, suggesting that the presence of fibres in the gut, and consequently the formation of SCFAs leading to lower pH, might attenuate microbial proteolysis. Indeed, it has been shown that p-cresol sulphate and phenylacetylglutamine were significantly lower in vegetarians than non-vegetarians^18^ and following a diet high in resistant starch^57^ while decreasing their levels via low-protein diets has been challenging^7^. Furthermore, supplementation with probiotics and/or prebiotics has been shown to decrease serum levels of p-cresol sulphate in chronic kidney disease patients^58^ and urinary levels in healthy volunteers^59^. Fibre availability in the colon therefore seems to play a role in the microbial fermentation of proteins that has been linked to unfavourable health outcomes^11,12^ and further investigations are required to disentangle the underlying mechanisms.

Although we recognize that our study is limited in cohort size and based on correlations and associations, it is to our knowledge the first study to link intestinal segmental transit times and pH with intra- and inter-individual differences in the gut microbiome composition and metabolism in a healthy population. While this study included a rather homogenous healthy group of volunteers (residents in Denmark, predominantly women), it provides valuable insights into longitudinal changes of faecal SCFAs and pH over a period of more than one week, which have not previously been documented. Our results show an important role of gut transit time and pH with regard to the inter-individual gut microbiome composition and production of microbial-derived metabolites. These results emphasise that the gut environment is important to consider in human microbiome studies in the quest for understanding the healthy gut microbiome and disentangling personal microbiome responses to foods and other lifestyle factors.

## Methods

### Study participants

A 9-day human study (PRIMA) among healthy subjects was conducted at the Department of Nutrition, Exercise and Sports (NEXS) at the University of Copenhagen in Denmark from April to December 2021. The research protocol was approved by the Municipal Ethical Committee of the Capital Region of Denmark (H-20074067) and all participants provided written informed consent to participation. The study was registered at ClinicalTrails.gov (ID: NCT04804319).

Sixty-one healthy participants living in Denmark (43 women and 18 men) were enrolled and completed the study. Participants were healthy by self-report (did not suffer from inflammatory bowel syndrome, small intestinal overgrowth, inflammatory bowel disease, chronic or infections disease, diabetes or cancer), aged 18-75 years with a BMI between 18.5 and 30.0 kg/m^2^ with no intake of medication with the exception of mild antidepressants and contraceptive pills. Intake of antibiotics, diarrhoea inhibitors and laxatives one month prior to the trial was not allowed. Furthermore, pregnant or lactating women were not included in the trial.

### Experimental design and sample collection

Seven days prior to the study, the participants were asked not to consume any sweet corn as two self-administered sweet-corn tests to evaluate the whole gut transit time were part of the study. Prior to both of the visits, the participants were asked to abstain from alcohol intake, smoking, and strenuous exercise.

The participants were asked to maintain their habitual diet and register their food intake online via the Myfood24 tool (myfood24.org) with nutritional values based on the Danish food composition database FRIDA version 4.1 (frida.fooddata.dk) for eight consecutive days during the study. During the trial, the participants collected daily stool samples (first bowel movement whenever possible), stored the samples in their domestic freezers and transported them to the laboratory while being kept cold. Moreover, the participants self-reported daily their defecation patterns including stool consistency assessed by the BSS and stool frequency, their physical activity, intake of dietary supplements and medication (limited to pain killers in a few cases), as well as their gastrointestinal symptoms. The gastrointestinal symptoms were assessed based on a 10 scale scoring system (0 – no symptoms, 10 – the most severe symptoms) in regards to stomach ache, bloating, constipation, diarrhea, and overall comfort. Furthermore, the participants collected seven daily spot morning urine samples (days 1, 2, 4, 5, 6, 7, 8) and two 24-hr urine samples (days 2-3 and days 8-9) during the study period. The collected urine samples were stored in participants’ domestic freezers, transported to the study site in a cooling bag, and stored at −20°C overnight. After thawing at 5°C, aliquots of 1 mL were taken and stored at - 80°C until further use. In addition, the participants consumed 100 g of sweet corn prior to their evening meal on days 3 and 5 and recorded the time of the corn egestion^17^.

At both of the visits (day 2 and day 9), fasting blood and breath samples were collected. During the first visit, anthropometric measurements (height, body weight, and BMI) were obtained. Furthermore, the first visit also included a standardized meal test for all participants (n = 61). The test meal consisted of rye bread (with butter and jam), a boiled egg, a portion of natural yoghurt along with nuts walnuts and blueberries, and a glass of water (100 ml) with 250 mg of dissolved paracetamol (**Table S1**), which was used as a marker of postprandial gastric emptying of liquids^60^. The meal portion size was calculated as 25 % of the daily energy demand of each participant based on the Harris-Benedict equation^22^. Postprandial urine samples (at 30 min, 60 min, 120 min, 180 min, 240 min, 300 min, 360 min, and between 6-8 h, 8-10 h, and 10-24 h) and postprandial breath exhalations (at 30 min, 60 min, 90 min, 120 min, 150 min, 180 min, 210 min, 240 min, 270 min, 300 min, 330 min, and 360 min) were collected. A subset of participants (n = 50) ingested a SmartPill® capsule immediately after the meal with a bit of additional water if needed. All participants drank 150 ml of water at 2 h and 4h after the meal, respectively. At 6 h, all participants received a sandwich and 500 ml of water and left the study site.

### SmartPill data collection and analysis

The SmartPill® capsule is a single-use wireless gastrointestinal capsule (26.8 mm x 13 mm), which transmits data on luminal pH, temperature, and pressure to a portable receiver, which was worn by the participants from ingestion to egestion and thereafter returned to the study personnel. The capsule measures a pH range of 1-9, with an accuracy of+-0.5 pH units, pressure at a range of 0-350 mmHg (± 5 mmHg), and temperature ranging between 20°C and 40 °C (±1 °C)^23^. Upon receiving the portable receiver, the raw data were downloaded from the receiver to the manufacturer’s software via a docking station. Intestinal segmental transit times were determined based on landmark changes in the pH values as follows: gastric emptying (GE) was defined as the time point with an abrupt increase of ≥ 3 pH units indicating passage from the stomach into the duodenum. The passage from the small intestine into the ileocaecal junction (ICJ) was defined as the first time point with a decrease of at least one pH unit. The body exit of the capsule was defined as the time point with a decrease in temperature and/or a loss of data. The time of capsule residence in each of the gastrointestinal segments corresponds to gastric emptying time (GET), small intestinal transit time (SITT), colonic transit time (CTT) and combined, whole gut transit time (WGTT). Regional pH and pressure profiles were also obtained and the median values were determined. The segmental transit time and pH values in the colon were further segmented into proximal, distal and recto-sigmoid, respectively. The proximal colon pH and transit time were estimated as median values of the first 32.3% of the total CTT, while for pH in the distal colon, median values of the next 32.6% of the total CTT, and for the recto-sigmoid pH the median pH of the last 35.4% of the total CTT was used based on previously reported data, which determined the percentages of total CTT according to the location of radio-opaque markers (visualized by X-rays) in the different segments of the colon^16^. In addition, the median pH value measured during the last 10 min prior to the capsule egestion was registered as rectal pH.

### Dietary records

Detailed 24-h weighted food intakes were recorded for 8 consecutive days by the participants via the online Myfood24 tool (myfood24.org) with nutritional values based on the Danish food composition database FRIDA version 4.1 (frida.fooddata.dk). The collected data included information about the intake of macronutrients (carbohydrate, protein, fat), dietary fibre (AOACFIB), coffee and alcohol intake in addition to information about more than 80 nutrients. Under-reporting was identified by calculating the average daily energy demand for each person divided by the reported caloric intake with a cut-off value of 0.8^61^. Accordingly, approximately 25 % of the daily dietary records were under-reported and the data were removed in the subsequent analyses in this study. In contrast, no over-reporters (> 2.5) were detected. The complete dietary profiles were used in the principal component analysis, whereas macronutrient profiles, coffee and alcohol intake were used in the redundancy analyses.

### Breath exhalations measurements

Fasting and postprandial levels of hydrogen and methane were measured in all breath samples by the M.E.C. Lactotest 202 Xtend device (M.E.C. R&D sprl, Brussels, Belgium).

### Biochemical analysis of blood

Blood samples were upon collection immediately put on ice until they were centrifuged for precipitation of blood cells and stored at −80°C. Glucose was measured in plasma samples by using Pentra ABX 400 (HORIBA ABX, Montpellier, France) with a detection limit of 0.11 mmol/L. Serum insulin and C-peptide levels were measured by using Immulite 2000 XPi (Siemens Healthcare Diagnostics Ltd., Llaneris Gwynedd LL554EL, UK) with the detection limit of 14.4 pmol/L and 27 pmol/L, respectively. Prior to the analyses, both instruments’ performances were validated using external and internal insulin, c-peptide and glucose controls. Three participants arrived for the second visit in a postprandial state, the blood was collected and analysed accordingly but the glucose, insulin and c-peptide values were not included in the data analysis.

### Faecal measurements

Faecal samples were upon receipt stored at −20°C overnight, thawed and homogenized in sterile water with a sample to water ratio of 1:1 (w/v) (faecal slurry). Subsequently, pH was measured in the faecal slurry using a digital pH meter (Mettler Toledo). The homogenized samples were subsequently aliquoted to cryotubes and stored at −80 °C until further analyses. Stool moisture was determined by evaporating the water of one aliquot (1 approximately 1 mL) using a vacuum concentrator (Speed-Vac, Christ RVC 2-25) and by calculating the faecal weight difference before and post-evaporation.

Faecal SCFAs and BCFAs were quantified by LC-MS in samples collected between day 2 and day 5 (n = 170) as previously described^34^. In brief, the samples were thawed, mixed with ethanol and purified by filtration (0.2 µm filter). Subsequently, the samples were derivatized with 3-nitrophenylhydrazine and labelled internal SCFA standards were added. Dilution series of external SCFA standards spiked with internal SCFA standards, and all derivatized samples were analyzed on UPLC-QTOF-MS (Synapt G2, Waters®) in negative ionization mode (cone voltage 3.0 kV) with an ACQUITY BEH C18 guard column (2.1 x 5 mm, 1.7 µm, Waters) coupled to an ACQUITY BEH C18 column (2.1 x 100 mm, 1.7 µm, Waters®) and with the collision energy of 6.0 eV. The faecal concentrations of SCFAs and BCFAs were determined using vendor software (Quanlynx, Waters®).

Bacterial load in faeces was determined using approximately 500 µL of frozen faecal slurry (238 – 816 mg) and diluting it 400,000 times in physiological solution (8.5 g/L NaCl; VWR International). Next, 1 ml of the microbial cell suspension obtained was stained with 1 μL SYBR Green I (1:100 dilution in dimethylsulfoxide; shaded 20 min incubation at 37 °C; 10,000 concentrate, Thermo Fisher Scientific). The flow cytometry analysis of the bacterial cells present in the suspension was performed using a Cytoflex flow cytometer (CytoFLEX 3; Beckman) as previously described (**Supplementary Fig. 8**)^18^. The final microbial load was calculated per gram of faeces.

### Microbiome profiling

DNA was extracted in random order from the faecal slurries (n=484) using DNeasy PowerLyzer PowerSoil kit (Qiagen, 12855-100) and the V3-region of the 16S rRNA gene was PCR amplified using 0.2 µl Phusion High-Fidelity DNA polymerase (ThermoFisher Scientific, F-553L), 4 µl HF-buffer, 0.4 µl dNTP (10 mM of each base), 1 µM forward primer (PBU; 5’-A-adapter-TCAG-barcode-CCTACGGGAGGCAGCAG-3’) and 1 µM reverse primer (PBR; 5’-trP1-adapter-ATTACCGCGGCTGCTGG-3’) and 0.05-5 ng faecal DNA in 20 µl total reaction volume. Both primers (TAG Copenhagen A/S) were linked to sequencing adaptors and the forward primer additionally contained a unique 10 bp barcode (Ion Xpress™ Barcode Adapters) for each sample. The PCR program consisted of an initial denaturation for 30s at 98 °C, followed by 24 cycles of 98 °C for 15 s and 72°C for 30 s, and a final extension at 72 °C for 5 min. The PCR products were purified by the HighPrep ™ PCR clean-up system (AC-60500 Magbio) according to the manufacturer’s protocol. The resulting DNA concentrations were determined by Qubit HS assay and libraries constructed with mixing equimolar amounts of each PCR product. Partial 16S rRNA gene sequencing was performed on an Ion S5™ System (ThermoFisher Scientific) using OneTouch 2 Ion5: 520/530 kit - OT2 400bp and an Ion 520 Chip. The raw data were pre-processed into an ASV table using our in-house pipeline^62^ based on the DADA2 algorithm and settings recommended for IonTorrent reads^63^, with taxonomy assigned to the ASVs using the RDP database (v18). The resulting ASV table, taxonomy and ASV sequences were merged into a phyloseq object for further analysis. For quantitative microbiome profiling (QMP) analyses, the relative abundances derived from the pre-processed 16S rRNA sequencing analysis were adjusted for the bacterial loads as previously published^64^. In brief, samples with < 10 000 reads were removed (n = 362) and downsized to even sampling depth, defined as the ratio between sample size (16S rRNA gene copy number corrected sequencing depth) and bacterial load. 16S rRNA gene copy numbers were retrieved from the ribosomal RNA operon copy number database rrnDB73^65^. The copy number corrected sequencing depth of each sample was rarefied to the level necessary to equate the minimum observed sampling depth in the cohort while assuring a minimum number of 10 000 reads in each sample and optimizing the chosen sampling depth to exclude as few samples as possible. In case of no copy number correction, an average copy number of 3.88 was used^6^.

### Metabolic profiling

#### Preparation of urine and faecal samples

Untargeted urine and faecal metabolomics were performed as previously published^34^. All urine samples were thawed on ice, centrifuged at 10,000*g* at 4 °C for 2 min, and transferred to a new tube to remove solid particles. The urine samples were kept cold on ice during preparation. Samples were randomized and pipetted into 15 plates (96-well). All urine samples from the same individual were placed on the same 96-well plate. Subsequently, they were diluted 1:5 with an internal standard mixture (L-Adenine-8-^13^C (Cambridge Isotope Lab), L-Phenyl-d5-Alanine-2,3,3-d3 (Cambridge Isotope Lab), Caffeic Acid ^13^C_3_ (Toronto Research Chemicals), Caffeine ^13^C_3_ (Toronto Research Chemicals), L-Tyrosine ^13^C_9_ (Sigma Aldrich), Para-aminobenzoic acid (Sigma Aldrich), L-Tryptophan- (indole-d_5_) (Sigma Aldrich), Hippuric Acid-[^13^C_6_] (IsoSciences), Cortisone-d8 (Sigma Aldrich), and Glycocholic Acid- [^2^H_4_] (IsoSciences). Quality control (QC) samples were obtained by mixing 20 µl of each urine sample in a given plate (plate pools) and by mixing 20 µl of each plate pool to create the global pool. The QC samples, blank assays (0.1% formic acid), and mixtures of known standards (including 33 microbial-derived compounds) were included in each plate. The plates were sealed and stored at 4 °C until analysis (24 h max, otherwise stored at −80 °C). If the plate was frozen and thawed again before analysis, the plate was gently mixed by vortex stirring for 30 min immediately prior to analysis.

Faecal homogenates collected between day 2 and day 5 (n = 170) were thawed at room temperature for 30 min and vortexed. Approximately 50 mg±5mg (≈50 µL) of the homogenates were mixed with 96 % ethanol, internal standard mixture (L-Adenine-8-^13^C (Cambridge Isotope Lab), L-Phenyl-d5-Alanine-2,3,3-d3 (Cambridge Isotope Lab), Caffeic Acid ^13^C_3_ (Toronto Research Chemicals), Caffeine ^13^C_3_ (Toronto Research Chemicals), L-Tyrosine ^13^C_9_ (Sigma Aldrich), Lysophosphatidylcholine (17:1d_7_) (Avanti Polar Lipids), L-Tryptophan-(indole-d_5_) (Sigma Aldrich), Hippuric Acid-[^13^C_6_] (IsoSciences), Cortisone-d8 (Sigma Aldrich), and Glycocholic Acid-[^2^H_4_] (IsoSciences)) resulting in a 1:60 dilution. The diluted samples were vortexed for 30 s and subsequently mixed at 60 °C for 2 min in a Thermo mixer at 1400 rpm, before being centrifuged at 14000 rpm (Eppendorf centrifuge 5417R), 4 °C for 2 min. The supernatants were filtered through a 0.2 µm filter and 200 µL of each faecal suspension was transferred to a 96-well plate, evaporated using a cooled vacuum centrifuge, and re-dissolved in 200 µL 0,1% formic acid prior to the UPLC-MS. All faecal samples from the same individual were placed on the same 96-well plate and QC samples were prepared in the same way as for the urine samples. In addition, each 96-well plate contained blank assays (96% ethanol) and mixtures of known standards (including 33 microbial-derived compounds).

#### UPLC-ESI-Q-TOF-MS Analysis

Both urine and faecal samples were profiled by ultra-performance liquid chromatography (UPLC) coupled with a quadrupole-Time of Flight Mass Spectrometer (q-TOF-MS) equipped with electrospray ionization (ESI) (Synapt G2, Waters®) in both positive and negative ionization mode^34^. Blank samples (0.1% formic acid), assay blanks, standard mixtures, and QC samples were injected regularly to evaluate LC-MS system stability, possible contamination and/or loss of metabolites. The injected samples (5 µL) were separated on a reversed-phase column (ACQUITY HSS T3 C18 column, 2.1×100 mm, 1.8 µm, Milford, USA) coupled with a pre-column (ACQUITY VanGuard HSS T3 C18 column, 2.1×5 mm, 1.8 µm, Milford, USA). The mobile phases consisted of 0.1% formic acid in water (solvent A) and 0.1% formic acid in 70:30 acetonitrile: methanol (solvent B). The duration of the analytical run was 7 min with the following flow rate: start condition (0.5 mL/min), 1 min (0.5 mL/min), 2 min (0.6 mL/min), 3 min (0.7 mL/min), 4 min (0.8 mL/min), 4.5 min (1.0 mL/min), 6.4 min (1.1 mL/min), 6.6 min (1.0 mL/min), 6.8 min (0.5 mL/min), 7.0 min (0.5 mL/min), and the following gradient: start condition (5% B), 1 min (8% B), 2 min (15% B), 3 min (40 % B), 4 min (70 % B), 4.5 min (100 % B), 6.6 min (5% B), 7 min (5% B). Mass spectrometry data were acquired in full scan mode with a scan range of 50-1000 mass/charge (*m/z*). Data-dependent acquisition (DDA) was performed on the top 3 most abundant ions on QC samples (only urine) to provide MS^2^ data. Electrospray settings were the following: the cone voltage was 2.5 kV and 3.2 kV; the collision energy was 6.0 and 4.0 eV, the temperature of the ion source and desolvation nitrogen gas temperature was 120 °C and 400 °C for positive and negative ionization mode, respectively.

#### Metabolite identification and structure elucidation

Tandem mass spectrometry (MS^2^) analyses were performed by a UHPLC system coupled to a Vion IMS QTOF mass spectrometer (Waters®) for obtaining spectra with higher mass accuracy. The samples were separated on a reversed-phase column (ACQUITY HSS T3 C18 column, 2.1×100 mm, 1.8 µm, Milford, USA) coupled with a pre-column (ACQUITY VanGuard HSS T3 C18 column, 2.1×5 mm, 1.8 µm, Milford, USA) at a temperature of 50 ℃. The mobile phases consisted of 0.1% formic acid in water (solvent A), methanol (solvent B), 0.1% formic acid in 70:30 acetonitrile: methanol (solvent C), and isopropanol (solvent D). The duration of the analytical run was 10 min with the following flow rate: start condition (0.4 mL/min), 0.75 min (0.4 mL/min), 6 min (0.5 mL/min), 6.5 min (0.5 mL/min), 8 min (0.6 mL/min), 8.1 min (0.4 mL/min), 9 min (0.4 mL/min), 10 min (0.4mL/min), and the following gradient: start condition (100% A), 0.75 min (100% A), 6 min (100% B), 6.5 min (70% C, 30 % D), 8 min (70% C, 30 % D), 8.1 min (70% C, 30 % D), 9 min (100% A), 10 min (100% A). Full scan acquisition was performed on selected urine samples with a scan range of 50-1500 *m/z*. Targeted MS^2^ was performed on a selected list of precursors at three different collision dissociation energies 10, 30, and 50 eV.

Mass spectra were manually interpreted and metabolites were identified by matching the precursor ion and fragmentation patterns with databases such as HMDB (https://hmdb.ca/), Metline (https://metlin.scripps.edu/), mzCloud (https://www.mzcloud.org/) and an in-house database. Furthermore, authentic standards were run together with the samples with the highest intensity on the same batch and platform. If needed, the authentic standards were sulfated or glucuronidated with either biomimetic synthesis^66^ or chemical synthesis^34^. The identification level of metabolites that were identified was classified according to Sumner *et al.* as level I (confirmed by matching to a standard with two orthogonal measures (rt, *m/z*), level II (matching MS^2^ fragmentation to a spectral library), level III (compound classification), or level IV (unknown)^25^. See **Supplementary Table 6 and Table 7** for further details. 3-Hydroxy-2-oxindole, 5-hydroxyoxindole, 2-picolinic acid, 4-methylcatechol, xanthine, 2-oxindole-3-acetic acid, pantothenic acid, nicotinic acid, tryptophan, sebacic acid, pipecolic acid, glutaric acid, citric acid, psedouridine, taurine, 1,3-dimethyluric acid, suberic acid, 1,3,7-trimethyluric acid were purchased from Sigma-Aldrich. 4-Hydroxyhippuric acid, 1-methylxanthine and 1-methyluric acid were purchased from Toronto Research Chemicals.

#### Metabolomics data processing

The raw data obtained by UPLC-MS were converted to mzML format by publicly available msConvert (ProteoWizard Toolkit)^67^. The converted data were pre-processed using the open-source R package XCMS (v3.18) using the centWave algorithm (requiring 3 consecutive scans with an intensity of over 10 counts)^68^. The pre-processing steps included noise filtering, peak picking, retention time alignment and feature grouping across samples, and filling of missing features, which were done separately for the urine and faecal samples (and for positive and negative mode), respectively. The detailed pre-processing parameter settings can be found in **Supplementary Table 8.** Noise filtering settings included that features should be detected in a minimum of 10 % of all samples. Features with a retention time below 0.5 min or above 6.8 min were excluded. Data tables were generated comprising mass-to-charge ratio (m/z), retention time (rt), and intensity (peak area) for each variable in every sample. Each detected peak is represented by a feature defined by a rt and a m/z. The obtained data were corrected for within- and between-batch intensity drift using the LOESS correction method^69^. The processed data were normalized by the probabilistic quotient normalization (PQN)^70^ method to correct for variations in urine and faecal concentrations within- and between-batches. Upon analyses of 15 plates with urine samples, QC samples clustered closely together in the principal component analysis (PCA) score plots confirming a stable UPLC system during the course of analysis with the exception of two plates in the negative mode and one plate in the positive mode, which had to be removed from further statistical analyses (**Supplementary Fig. 9**).

Moreover, features with high variability after normalization across the pooled QC samples were filtered out (CV% > 50 %). Finally, the CAMERA package^71^ (v1.52) was used to group features together based on retention time (tolerance = 0.1s) and to annotate possible adducts and isotopes.

#### Statistical Analysis

Statistical analyses were conducted in R (v 4.2). The area under the curves (AUC) for hydrogen and methane concentrations during the postprandial period was calculated using the trapezoid rule in GraphPad Prism (v 9.2.0). The normality of data was assessed with the Gaussian distribution and Shapiro-Wilk test procedure.

Mixed-effects linear regression models were used to examine the day-to-day fluctuations and inter-individual variation in gut environmental factors using data from all 9 days. The models were generated using the *lme4* R package (v 1.1-31) as lmer(gut environmental factor ∼ factor(Day) + (1 | Participant ID), moreover ranova function from the *lmerTest* package (v 3.1-3) was used to perform the random effects-likelihood ratio tests to infer whether Participant ID significantly contributes to explaining the variation in the gut environmental factors. A p-value of < 0.05 was considered statistically significant. Coefficients of intra-individual variation were calculated as CV_intra_ = (SD_intra_ / Mean_intra_) * 100 where mean and SD were based on all measurements from a single individual over the 9 days.

Gut microbiome beta-diversity analysis using Bray Curtis distances as well as metabolome and diet beta-diversity analyses using Euclidian distances were performed with the *phyloseq* package (v 1.42.0) and PERMANOVA tests by adonis2 function from the *vegan* package (v 2.6) with 999 permutations and strata = Participant ID when testing the day-to-day fluctuations.

Single time point correlations were calculated using standard Spearman’s rank correlation, as implemented in the *Hmisc* R package (v 4.7), and heatmaps were generated by the *corrplot* package (v0.92). Repeated measure correlations were performed using the *rmcorr* (v 0.5)^72^.

Distance-based redundancy analysis (db-RDA) was performed to quantify the effect sizes of gut environmental factors and other variables on the intra-individual and inter-individual variation in the gut microbiome (both relative and quantitative profiles at genus level), faecal metabolome, and urine metabolome. The analyses were performed with Bray-Curtis dissimilarity using the *capscale* function as implemented in the *vegan* package (v 2.6). With regards to intra-individual analyses, data available from all samples (day 1-day 9) and *strata* = Participant ID were used. For the inter-individual analyses, data collected on day 2 (visit 1) were used separately for all participants (n = 61) and for the SmartPill subgroup (n = 50). The statistical significance was determined by permutation test with 9999 random permutations (*anova.cca* function) and p-values were adjusted for multiple testing by false discovery rate (Benjamin–Hochberg)^73^, an adjusted p-value (q-value) below 0.1 was considered significant.

For the untargeted metabolomics data, the area of each m/z feature was log-transformed and missing values were imputed and replaced by values reflecting half of the minimum intensity of the given m/z feature. Linear regression models and sparse partial least squares (SPLS) models were performed to examine the relationship between the m/z features and the variables of interest (i.e. segmental transit time and pH). The modelling was performed using the SmartPill-derived data and the 24-h postprandial urine metabolome collected at day 2 as well as the faecal metabolome closest to the time of the SmartPill egestion. The linear mixed models were performed with the *lme4* R package (v 1.1-31). The multivariate SPLS models were performed with the *caret* R package (v 6.0-92). P values were corrected for multiple testing by the Benjamin–Hochberg false discovery rate (q-value). Features with a q-value < 0.1 were considered to be statistically significant and only features selected by both the linear regression and SPLS were further submitted for identification including the MS^2^.

## Data availability

All sequencing data have been submitted to the NCBI Sequence Read Archive (SRA). BioProject ID: PRJNA1027590.

## Code availability

No custom code was generated for this work.

## Acknowledgements

We thank all of the dedicated volunteers that participated in the PRIMA study. We acknowledge our colleague Jan Stanstrup for help with the metabolomics data preprocessing, the students Anett Seer, Inès Barrau, Stine Hjort Kaas, and Kirstine Lea Lind Clemmensen, as well as the bioanalysts Sarah Fleischer Ben Soltane and Jane Guldborg Jørgensen for their assistance during the study visits and laboratory work at the Department of Nutrition Exercise and Sports. We would also like to thank Özge Cansin Zeki for her help with metabolite annotation. Furthermore, we thank Katja Ann Kristensen (Technical University of Denmark) for the preparation for sequencing. We also thank Duyen Nguyen (KU Leuven) for the help with flow cytometry and Jorge F. V. Castellanos (KU Leuven) for the help with QMP analysis. This study was supported by the Novo Nordisk Foundation (NNF19OC0056246; PRIMA—toward Personalized dietary Recommendations based on the Interaction between diet, Microbiome and Abiotic conditions in the gut). LOD was supported by a Semper Ardens grant from the Carlsberg Foundation. The funding bodies played no role in the design of the study, collection, analysis, and interpretation of data, or the writing of the manuscript.

## Author contributions

NP, TRL, LOD and HMR conceived and designed the human study as part of the PRIMA collaboration headed by TRL. NP conducted the study under the supervision of LOD and HMR. Urine metabolomics was performed by NP. Metabolite annotations were done by NP, GLB and LOD. Metabolite synthesis and fine identification were done by GLB. Faecal SCFAs were analysed by ET and NP. The faecal metabolome was analysed by MSJ and NP. MFL generated the microbiome data. Bacterial load was done by NP under the supervision of JR. Statistical analyses were performed by NP with help from MFL and MAR. Expert supervision was performed by JR, TRL, LOD, and HMR. NP and HMR drafted the manuscript. All authors contributed to and approved the final manuscript.

