## Supplemental files for "Gut environmental factors explain variations in the gut microbiome composition and metabolism within and between healthy adults"

| **Supplementary Table 1. Energy and macronutrient composition of the standardized meal test provided at visit 1.** | | | | | |
| --- | --- | --- | --- | --- | --- |
| The final amounts were personalized corresponding to 25 % of the daily energy demand of each participant. | | | | | |
|  | **Amount (g)** | **Energy (kcal)** | **Carbohydrate (g)** | **Protein (g)** | **Fat (g)** |
| **Water** | 100 | 0 | 0 | 0 | 0 |
| **Rye bread** | 45 | 90.7 | 16.8 | 2.29 | 0.63 |
| **Butter** | 10 | 72.8 | 0.12 | 0.07 | 8.15 |
| **Strawberry jam** | 10 | 17.4 | 4.2 | 0.05 | 0 |
| **Egg (medium size)** | 50 | 68.2 | 0.55 | 6 | 4.7 |
| **Natural yoghurt** | 100 | 50.4 | 4.8 | 4.1 | 1.6 |
| **Blueberries** | 25 | 13.1 | 2.6 | 0.17 | 0.13 |
| **Walnuts** | 25 | 171 | 2.83 | 3.58 | 16.1 |
| **Total** | **265** | **484** | **31.9** | **16.3** | **31.3** |

**CONSORT Flow Diagram**

Analysed (n = 61)
♦ Excluded from analysis (urine metabolome data of 12 individuals were excluded due to large batch effect)

Lost to follow-up (give reasons) (n = 0)

Discontinued intervention (give reasons) (n = 0)

### Visit 1 and 2

Allocated to intervention (n = 63)

♦ Received allocated intervention (n = 61)

♦ Did not receive allocated intervention (give reasons) (n = 2, could not participate due illness and antibiotics exposure)

### Allocation

### Enrollment

Excluded (n = 28)

♦  Not meeting inclusion criteria (n = 21)

♦  Declined to participate (n = 7)

Enrolled (n = 63)

Assessed for eligibility (n = 91)

### Analysis

**Supplementary Figure 1.** CONSORT flow diagram of the PRIMA study.

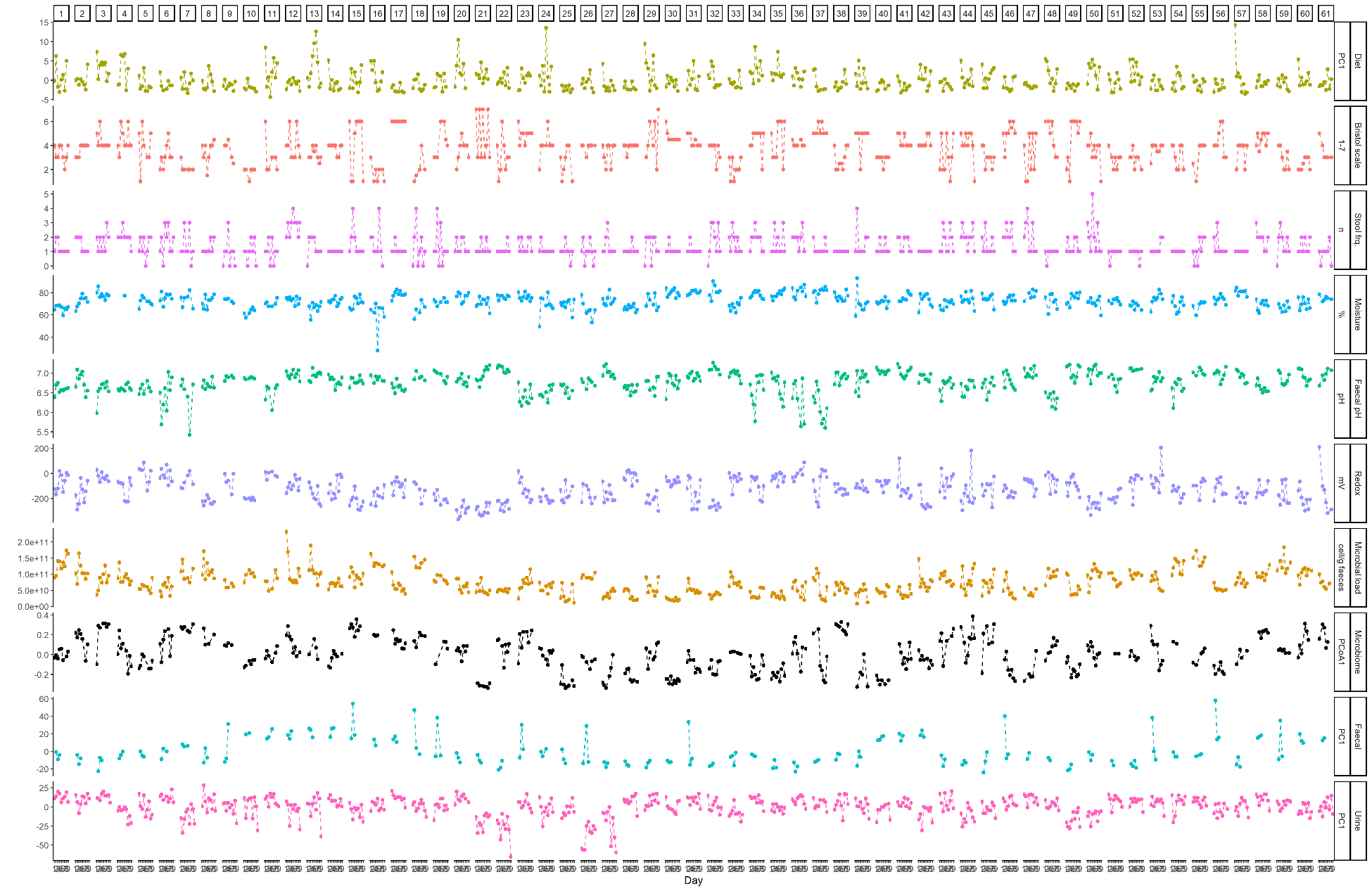

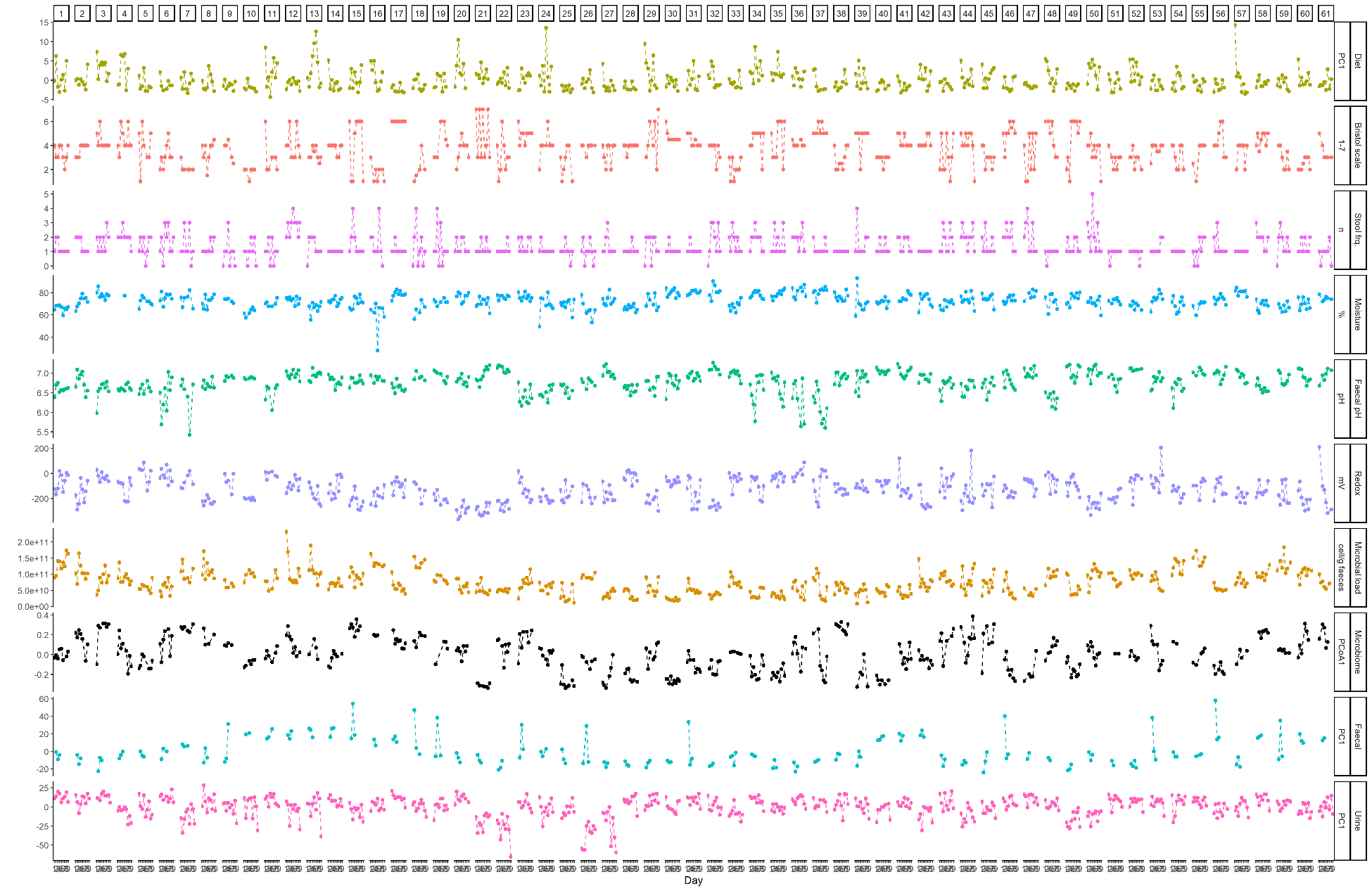
**Supplementary Figure 2.** Intra-individual fluctuations in diet, gut environmental factors, faecal microbiome, and urine and faecal metabolomes of all 61 individuals over the 9 days.

| **Supplementary Table 2. Mixed effect models to infer day-to-day and inter-individual variations in the gut environmental factors**  **A** | | | | | | | |
| --- | --- | --- | --- | --- | --- | --- | --- |
|  | **model1 = lmer(gut environmental factor ~ factor(Day) + (1 \| Participant ID))** | | | | | | |
| Gut environmental factor | term | sumsq | meansq | NumDF | DenDF | statistic | p.value |
| Stool moisture | factor(Day) | 391.95 | 48.99 | 8 | 406.69 | 2.17 | **0.029** ^a^ |
| Bristol scale | factor(Day) | 25.50 | 3.19 | 8 | 408.00 | 2.84 | **0.004** ^a^ |
| Faecal pH | factor(Day) | 0.25 | 0.03 | 8 | 408.02 | 0.81 | 0.596 ^b^ |
| Stool frequency | factor(Day) | 15.94 | 1.99 | 8 | 421.58 | 5.05 | **<10^-5^** ^a^ |
| Microbial load | factor(Day) | 9.23**⋅**10^21^ | 1.15**⋅**10^21^ | 8 | 23935589 | 2.56 | **0.009** ^a^ |
| **B** | **ranova(model1)** | |  |  |  |  |  |
| Gut environmental factor | term | effect | estimate | std.error | statistic | df | p.value |
| Faecal pH | Participant ID | fixed | 6.77 | 0.04 | 181.30 | 187.53 | **<10^-210^** |
| Bristol scale | Participant ID | fixed | 4.02 | 0.18 | 22.55 | 278.06 | **<10^-64^** |
| Stool frequency | Participant ID | fixed | 1.36 | 0.09 | 14.57 | 403.97 | **<10^-38^** |
| Moisture | Participant ID | fixed | 71.98 | 0.87 | 82.73 | 219.98 | **<10^-166^** |
| Cell count | Participant ID | fixed | 7.48E+10 | 4.8E+09 | 15.69 | 131.07 | **<10^-31^** |

^a^ p < 0.05 indicates that the gut environmental factor fluctuated from day-to-day, ^b^ p > 0.05 indicates that the factor was stable over days

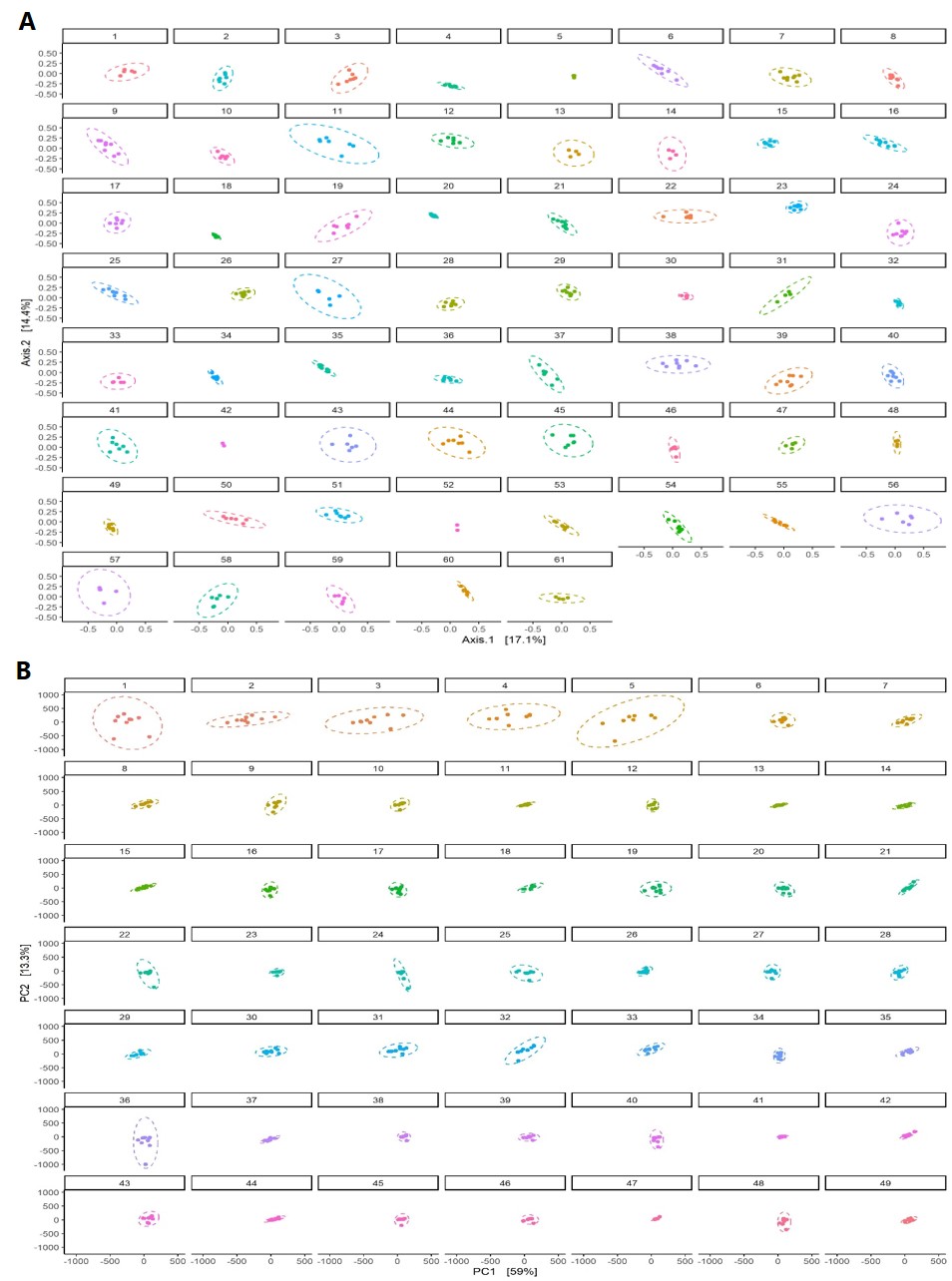

**Supplementary Figure 3.** **(A)** Individual Bray Curtis β-diversity ordination of quantitative faecal microbiome profiles over 9 study days (n=61) **(B)** Euclidian β-diversity ordination of urinary metabolomes profiles over 9 study days (n = 49, 12 individuals were removed due to LC-MS batch effect)

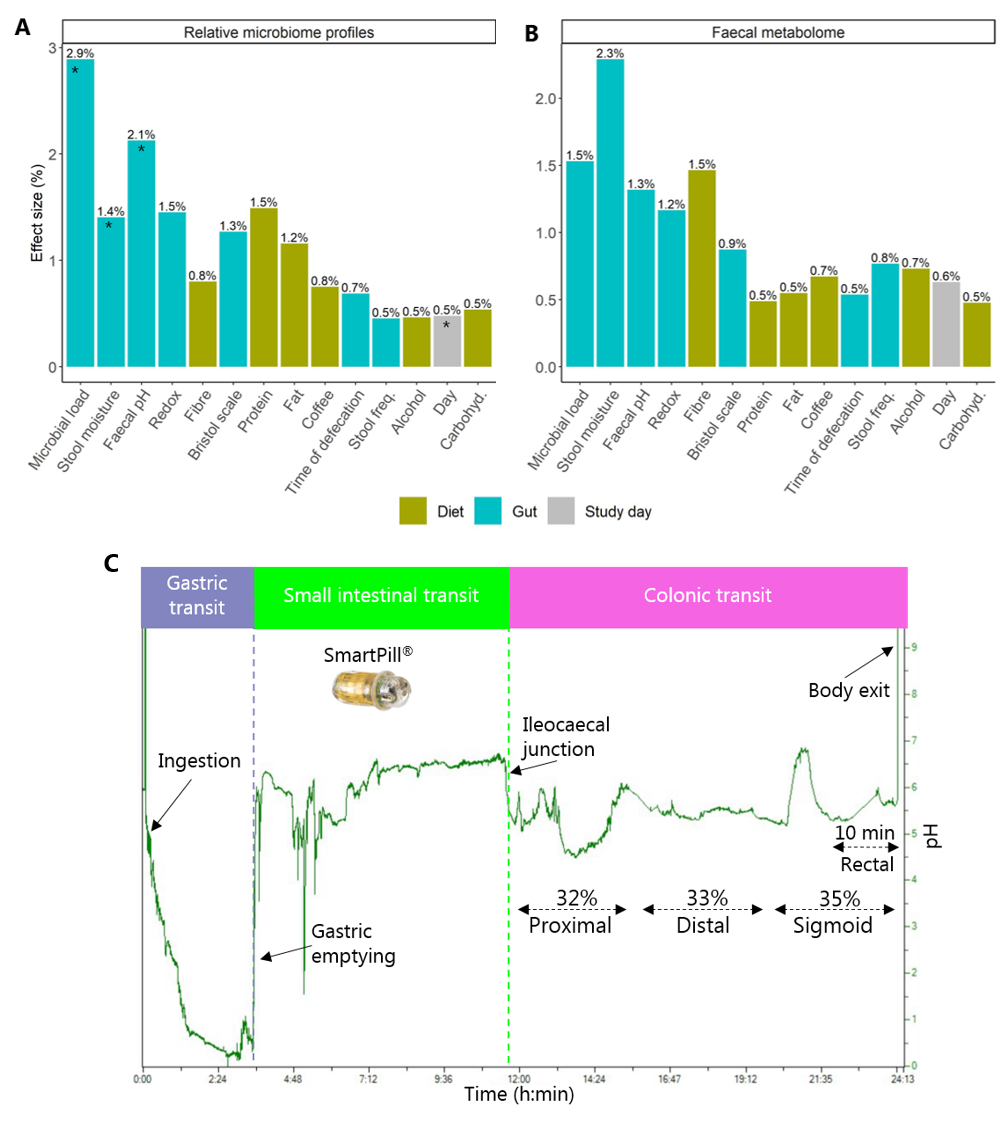

**Supplementary Figure 4. Contributions of dietary and gut factors on intra-individual variations in (A) relative microbiome profiles and (B) faecal metabolome.** The analysis was performed with distance-based redundancy analysis (db-RDA) with permutation test on daily relative microbiome data and untargeted faecal metabolome data. The asterisks indicate statistical significance (*q-value < 0.05). **(C) An example of a pH profile measured by the SmartPill.** Segmental transit times were determined based on pH changes upon gastric emptying, ileocaecal junction and body exit as indicated. The proximal-, distal-, and sigmoid-colon pH were determined as median values in each of the segments of the colon based on an approximation of the transit time based on previous data showing that the first 32% followed by 33 % and 35 % of CTT corresponds to the proximal, distal, and sigmoid colon, respectively.^74^ In addition, the median pH of 10 min before the capsule egestion was registered as rectal pH.

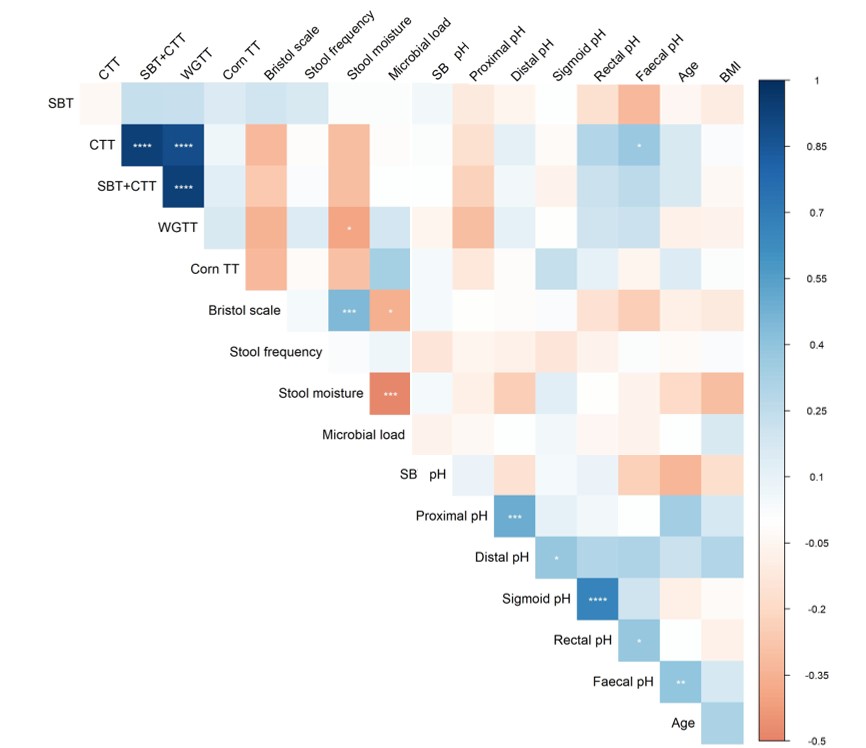

**Supplementary Figure 5. Spearman correlation analysis between segmental transit times assessed by the SmartPill, corn transit time, various proxy markers of transit time, gut factors, and subject characteristics.** The colour gradient shows the Spearman correlation coefficient and the asterisks indicate statistical significance (****q<0.001,***q<0.01,** q<0.05,*q<0.1). SBT; small bowel transit time, CTT; colonic transit time, WGTT; whole gut transit time).

Supplementary Tables 3 and 4 are enclosed as Excel files.

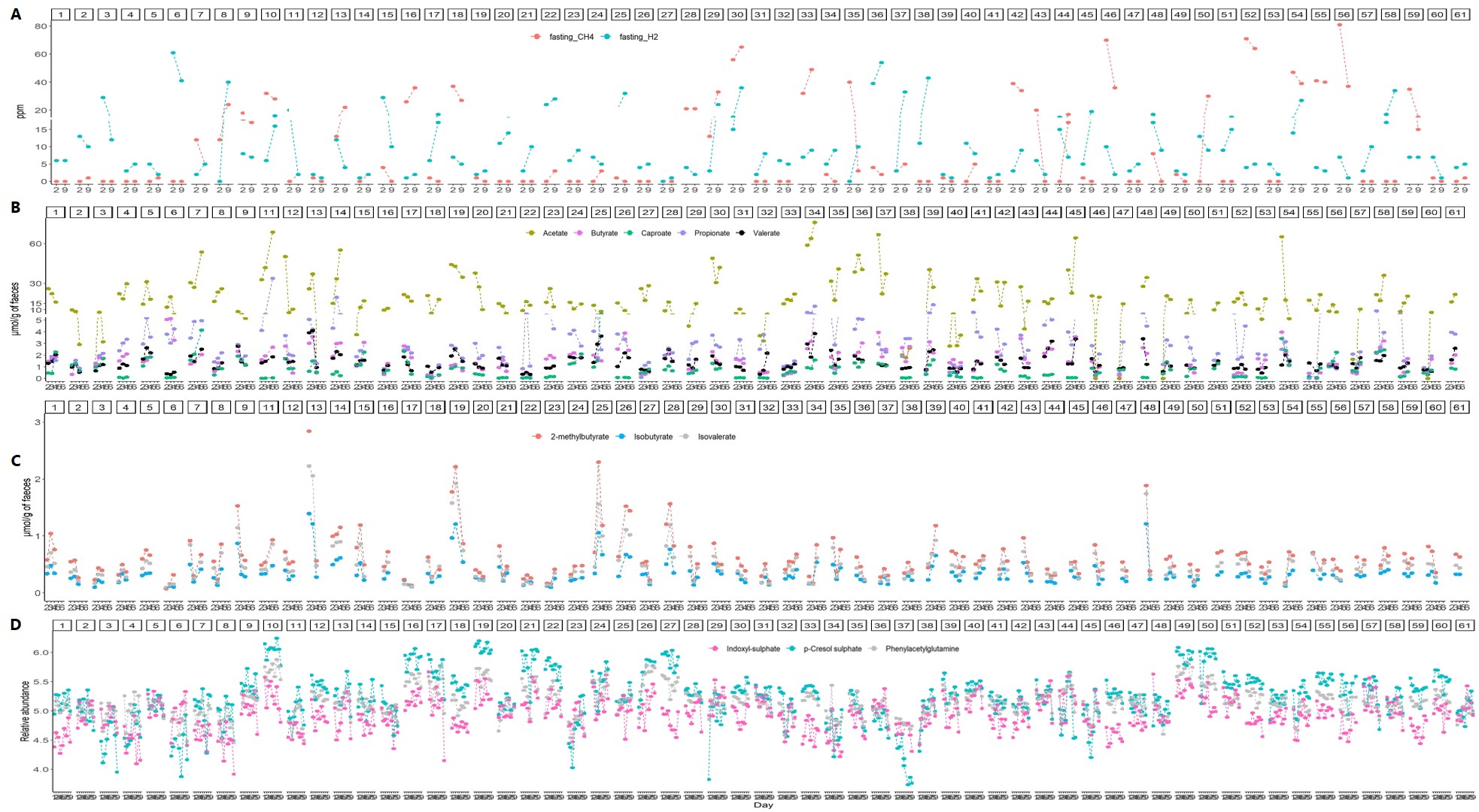
 **Supplementary Figure 6. Intra-individual variations in microbial-derived metabolites**. (A) Fasting breath hydrogen and methane levels (ppm) at days 2 and 9. (B) Concentrations of faecal short-chain fatty acids (SCFAs) and (B) branched chain-fatty acids (BCFAs) over 4 days in all 61 participants. (C) Relative abundances of urinary markers of microbial proteolysis over 9 days in all 61 participants

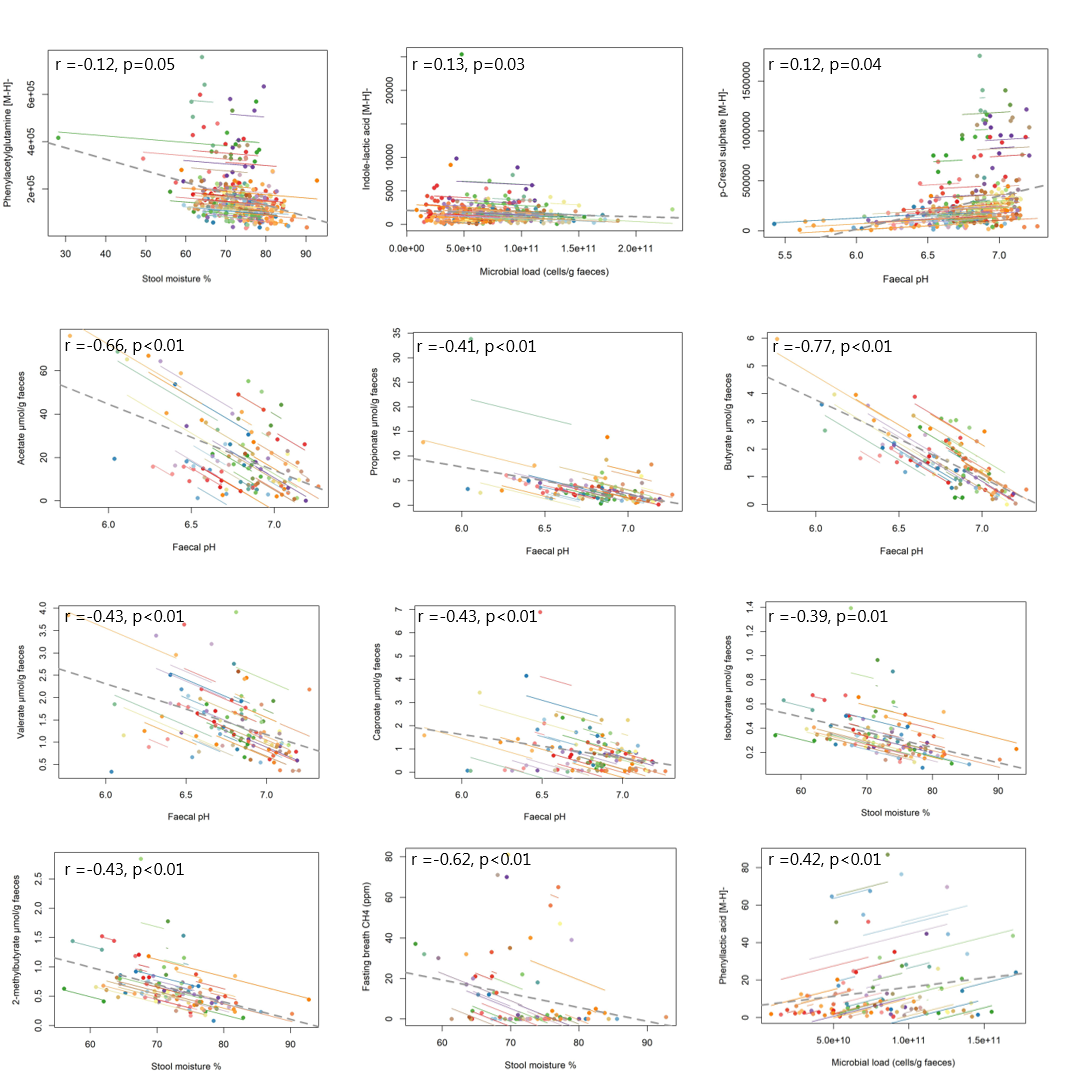

**Supplementary Figure 7. Repeated measures correlation between gut factors and microbial metabolites.** The colour lines show the individual correlation between each pair of tested variables for each of the study days using data from all 61 participants. The grey dashed lines, the correlation coefficients (r) and the p values indicate the overall trends.

**Supplementary Table 5. Repeated measurement correlations,** performed with rmcorr package

| **Dietary component** | **Metabolite** | **df** | **rmcorr.r** | **lowerCI** | **upperCI** | **p.val** | **effective.N** | **q.val** |
| --- | --- | --- | --- | --- | --- | --- | --- | --- |
| Protein | Hippurate | 303 | 0.17 | 0.06 | 0.28 | 0 | 305 | 0.02 |
| Alcohol | Propionate | 49 | 0.39 | 0.13 | 0.6 | 0 | 51 | 0.05 |
| Coffee | Indoxyl-glucuronide | 303 | -0.14 | -0.25 | -0.03 | 0.01 | 305 | 0.08 |
| Vegetables | TMAO | 303 | -0.14 | -0.25 | -0.03 | 0.01 | 305 | 0.08 |
| Starch | TMAO | 303 | -0.13 | -0.24 | -0.02 | 0.02 | 305 | 0.09 |
| Fibre | Phenylacetylglutamine | 303 | -0.12 | -0.23 | -0.01 | 0.03 | 305 | 0.09 |
| Fibre | p-Cresol sulphate | 303 | -0.12 | -0.23 | 0 | 0.04 | 305 | 0.09 |
| Fat | Tryptophan | 303 | -0.13 | -0.24 | -0.02 | 0.02 | 305 | 0.11 |
| Fibre | Isobutyrate | 49 | -0.32 | -0.54 | -0.04 | 0.02 | 51 | 0.14 |
| Fibre | Isovalerate | 49 | -0.31 | -0.54 | -0.04 | 0.03 | 51 | 0.14 |
| Fibre | Tryptophan (f) | 49 | 0.31 | 0.04 | 0.54 | 0.03 | 51 | 0.14 |
| Fat | Butyrate | 49 | -0.32 | -0.54 | -0.04 | 0.02 | 51 | 0.14 |
| Starch | Butyrate | 49 | -0.31 | -0.54 | -0.04 | 0.03 | 51 | 0.14 |
| Starch | Acetate | 49 | -0.31 | -0.54 | -0.03 | 0.03 | 51 | 0.14 |
| Protein | Indole-acetic acid | 303 | 0.12 | 0.01 | 0.23 | 0.03 | 305 | 0.16 |
| Carbohyd. | Acetate | 49 | -0.29 | -0.53 | -0.02 | 0.04 | 51 | 0.16 |
| Fibre | 2-methylbutyrate | 49 | -0.29 | -0.52 | -0.01 | 0.04 | 51 | 0.18 |
| Fat | Indoxyl-glucuronide | 303 | -0.12 | -0.23 | -0.01 | 0.04 | 305 | 0.18 |
| Carbohyd. | Butyrate | 49 | -0.28 | -0.52 | 0 | 0.05 | 51 | 0.19 |

Only p < 0.05 are displayed, SCFAs (acetate, propionate, butyrate, caproate, valerate) and BCFAs (2-methylbutyrate, isovalerate, isobutyrate) were measured in faeces, other metabolites in urine unless indicated otherwise. (f); faecal metabolite, carbohyd.; carbohydrates

**Supplementary Table 6. Metabolites identified by LC-MS/MS associated with segmental transit time and/or pH**

| **Metabolite^a)^** | **RT** | ***m/z*** | **ppm** | **Annotation** | **Adducts** | **MS^2^ fragments** | **Association** | **Regression coefficient** | **p-value^b)^** |
| --- | --- | --- | --- | --- | --- | --- | --- | --- | --- |
| **Urine** | | | | | | | | | |
| 4-Hydroxybenzoic acid sulphate^II^ | 3.89 | 216.9806* | 1 | [M-H]^-^ |  | 137.0241*, 124.0165, 109.0284,93.0344 | CTT | 0.04 | 0.0019 |
| 5-Hydroxy-2-oxindole sulphate^II^ | 3.53 | 227.9966* | 0 | [M-H]^-^ |  | 182.9546, 164.9429, 151.0399, 148.0399*, 145.0980, 138.9658, 136.9493, 120.0451, 107.0495, 105.0345, 92.0502, 79.9576, 68.0856 | WGTT/CTT | 0.04/0.04 | <0.0001/0.00008 |
| 3-Hydroxy-2-oxindole-sulphate^III^ | 4.02 | 227.9967* | 0 | [M-H] ^-^ | 228.9994 [M+1]^-^ 457.0000 [2M-H]^-^ 478.9850  [2M-2H+Na]^-^ | 186.0231, 150.0559, 148.0403*, 142.0329, 120.0450, 107.0380, 79.9569 | WGTT/Faecal pH | 0.03/1.25 | 0.0096/0.0399 |
| p-Cresol sulphate^I^ | 3.48 | 187.0063* | 1 | [M-H] ^-^ |  | 107.0494*, 79.9561* | WGTT/Sigmoid pH/Rectal pH/Faecal pH | 0.04/0.63/0.65/1.55 | 0.0009/0.0009/0.0003/<0.0001 |
| Phenylacetylglutamine^I^ | 3.21 | 265.1190* | 1 | [M+H]^+^ |  | 248.093*,147.0771*,130.0507*,101.0710*,84.0446 | WGTT/Sigmoid pH/Rectal pH/Faecal pH | 0.03/1.67/1.53/2.34 | 0.0010/0.0001/0.0003/<0.0001 |
| 1-Methylxanthine^I^ | 3.94 | 167.0555* | 8 | [M+H]^+^ |  | 110.0338, 82.0437, 55.0307 | SB pH | -1.64 | 0.0010 |
| 1,3-Dimethyluric acid^I^ | 4.18 | 195.0517* | 1 | [M-H]^-^ |  | 180.0290, 138.0300, 137.0230, 123.0080, 110.0360, 93.0331, 83.0253, 80.0257 | SB pH | -0.74 | 0.0006 |
| 1-Methyluric acid^I^ | 3.7 | 181.0357* | 3 | [M-H]^-^ | 363.083  [2M-H]^-^  385.061  [2M-2H+Na]^-^ | 138.0320, 122.0230, 110.0350, 96.0203, 83.0244, 69.0091 | SB pH | -1.69 | 0.0007 |
| Taurine^I^ | 0.66 | 124.0065* | 3 | [M-H]^-^ |  | 124.0065, 106.9801, 94.9800, 79.9569 | SB pH | 1.45 | 0.0012 |
| Pseudouridine^I^ | 1.12 | 243.0630 | 5 | [M-H]^-^ |  | 225.0503, 183.0405,153.0297*,140.0641, 110.0256 | Sigmoid pH | -0.50 | 0.0009 |
| Pimelic acid^I^ | 5.18 | 159.0658 | 0 | [M-H]^-^ |  | 141.0550, 115.0761*, 97.0655, 95.0489 | Rectal pH | -0.63 | 0.0002 |
| 4-Methylcatechol sulphate^II^ | 4.85 | 203.0016* | 1 | [M-H]^-^ |  | 203.0016, 123.0445*, 122.0369, 108.0216, 105.01343, 95.0505, 79.9871 | Rectal pH | 0.59 | 0.0008 |
| 4-Hydroxyhippuric acid sulphate^II^ | 3.86 | 274.0022* | 0 | [M-H]^-^ |  | 230.0122,194.0458, 150.0558, 93.0354 | SB pH | -1.69 | <0.0001 |
| 1,7-Dimethyluric acid^II^ | 4.58 | 195.0517* | 1 | [M-H]^-^ |  | 180.0286,162.8392, 160.8415, 138.0308, 137.0228, 123.0067, 121.9991, 110.0355, 93.0350, 67.0302 | SB pH | -0.55 | <0.0001 |
| 1,3,9-Trimethyluric acid^III^ | 4.88 | 211.0895* | 0 | [M+H]^+^ |  | 196.0666, 183.0949, 167.0993, 154.0652, 139.0418, 126.0709, 124.0935, 111.0440, 110.0385, 83.0602 (30eV) | SB pH | -0.72 | 0.0012 |
| Citric acid^I^ | 1.33 | 191.019 | 1 | [M-H]^-^ | 405.0278* [2M-H+Na]^-^ 383.0456  [2M-H]^-^ | 173.0090, 154.9980,129.0190, 111.0080, 87.0086, 85.0294,67.0188,57.0345 | Proximal pH | 1.42 | <0.0001 |
| Ascorbic acid glucuronide^III^ | 1.02 | 351.0569* | 1 | [M-H]^-^ |  | 333.0477, 291.0357, 241.9762, 235.0393, 193.0348, 175.0246, 162.0196, 157.0140, 115.0035, 87.0087 | Faecal pH | 2.89 | 0.007 |
| Argininic acid^III^ | 0.93 | 176.1020* | 8 | [M+H]^+^ |  | 159.0773, 134.0802, 130.0963, 112.0751, 84.0810, 71.0492, 70.0674, 60.0575 | Rectal pH | -0.75 | <0.0001 |
| Picolinoylglycine^III^ | 4.66 | 181.0600 | 7 | [M+H]^+^ |  | 163.0502*, 135.0544, 124.0489, 106.0276, 96.0431, 78.0325, 69.0439, 67.0281,55.0284,51.0199 | Sigmoid_pH/rectal pH | 0.7/0.7 | <0.0001/<0.0001 |
| Dihydroferulic acid glucuronide^III^ | 4.97 | 371.0981* | 1 | [M-H]^-^ |  | 353.0882, 326.1336, 291.0848, 195.066, 175.0246, 135.0445, 113.0244, 85.0293, 71.0128, 59.0145 | Rectal pH | -1.23 | 0.0008 |
| 3-Hydroxy-2-oxindole glucuronide^II^ | 4.5 | 324.0724* | 1 | [M-H]^-^ |  | 193.0351, 175.0246, 160.0402,148.0402, 145.0143, 131.0350, 113.0242, 103.0031, 101.0244, 89.0244, 85.0294, 72.9924, 71.0138, 59.0134 | Faecal pH | 1.12 | 0.0189 |
| **Faeces** | | | | | | | | | |
| Pantothenic acid^I^ | 4.19 | 220.1190* | 2 | [M+H]^+^ | 242.097  [M+Na]^+^  258.041  [M+K]^+^ | 202.1067, 184.0962, 142.0847, 124.0750, 116.0333, 98.0236, 90.0544, 72.0432, 67.0550 | CTT/Faecal pH | -0.04/-1.19 | 0.0031/<0.0001 |
| 2-Oxindole-3-acetic acid^I^ | 5.36 | 192.0655 | 3 | [M+H]^+^ | 214.045  [M+Na]^+^ | 192.0655, 174.0544, 146.0596*, 135.0433, 128.0493, 120.0800, 118.0648, 107.0748, 104.0491, 101.0378, 91.0552, 79.0554, 77.0385 | WGTT/CTT/Faecal pH | -0.1/-0.09/-1.81 | 0.0008/0.0014/<0.0001 |
| Pipecolic acid^I^ | 0.99 | 130.0871* | 2 | [M+H]^+^ |  | 112.0734, 84.0802, 56.0489, 55.0547 | WGTT/CTT | -0.08/-0.07 | 0.0008/0.0026 |
| Pimelic acid^I^ | 5.17 | 159.0658 | 0 | [M-H]^-^ |  | 159.0658, 141.0550, 115.0761, 97.0655*, 95.0489 | WGTT/CTT | 0.11/0.11 | 0.0012/0.0013 |
| 1-Methylxanthine^I^ | 3.91 | 165.0430* | 10 | [M-H]^-^ |  | 147.0404, 126.0330,108.0198, 80.0249, 65.9984 | WGTT/Faecal pH | -0.14/-3.03 | 0.0010/<0.0001 |
| Glutaric acid^I^ | 3.20 | 131.0359* | 11 | [M-H]^-^ | 301.0276  [2M+K-2H]^-^ | 87.0453*, 85.0295, 76.9828, 69.0351, 59.0142 | WGTT/Sigmoid pH/Rectal pH/Faecal pH | -0.09/-2.17/-1.84/  -2.56 | 0.0017/0.0003/0.0019/<0.0001 |
| Suberic acid^I^ | 5.72 | 173.0814 | 0 | [M-H]^-^ |  | 129.0912, 111.0814*, 102.9716, 86.9762, 83.0504 | WGTT | 0.04 | 0.0009 |
| Sebacic acid^I^ | 6.29 | 201.1123 | 2 | [M-H]^-^ |  | 183.1020*, 139.1124, 111.0826, 61.9879 | WGTT | 0.05 | 0.0012 |
| Cholic acid^III^ | 6.74 | 407.279 | 2 | [M-H]^-^ | 838.5542*  [2M-H+Na+1]^-^  837.543  [2M-H+Na]^-^815.566  [2M-H]^-^ 443.2573*  [M+Cl]^-^ | 407.2796, 390.9812, 386.9874, 366.9807, 350.9871, 345.2803, 342.9791, 340.9842, 324.9888, 304.9824, 302.9860, 290.9892, 270.9793, 264.9896, 254.9873, 242.9851, 240.9892, 204.9896, 192.9872, 191.9430, 174.9574, 154.9916, 146.9877, 130.9921, 92.566 | SB pH/Faecal pH | 2.00/-1.89 | 0.0010/<0.0001 |
| Unknown bile acid glycine conjugate^III^ | 6.78 | 470.3420* | #VALUE! | [M+H]^+^ | 492.3252  [M+Na]^+^  939.674  [2M+H]^+^ | 470.3422, 452.3318, 434.3215, 416.3105, 341.2781, 323.2611, 297.2496, 227.1424, 209.1315, 171.1384, 158.0811, 123.0807, 110.1054, 109.0653, 107.0845, 95.0856, 93.0686, 76.0404, 69.0731 | Distal pH | 0.60 | 0.0018 |
| Xanthine^I^ | 1.66 | 151.0270* | 9 | [M-H]^-^ | 152.0304*  [M+1]^-^ | 151.0254, 133.0147, 108.0197, 80.0263, 65.9971 | Sigmoid pH/Rectal pH | -1.12/-0.85 | <0.0001/0.0012 |
| Deoxyxanthosine^III^ | 4.04 | 267.0720* | 4 | [M-H]^-^ |  | 177.0404, 151.0259, 134.0360, 108.0200, 92.0254 | Rectal pH | -1.07 | 0.0015 |
| Tryptophan^I^ | 4.44 | 205.0983* | 3 | [M+H]^+^ |  | 188.0698*, 170.0592, 159.0906, 146.0593, 144.0797, 132.08155, 118.0649 | Faecal pH | -0.80 | 0.0003 |
| Nicotinic acid^I^ | 1.17 | 124.0402* | 3 | [M+H]^+^ |  | 106.0290, 98.9608, 96.0451, 84.9598, 80.0493, 78.0332 | Faecal pH | -0.99 | <0.0001 |
| Pseudouridine^I^ | 1.12 | 243.0615 | 1 | [M-H]^-^ |  | 225.0503, 183.0405,153.0297*,140.0641, 110.0256 | Faecal pH | -1.23 | <0.0001 |
| 4-Hydroxyphenyllactic acid^II^ | 2.62 | 181.0495* | 3 | [M-H]- |  | 163.0398, 135.044, 119.0493, 107.0496, 72.9947 | Faecal pH | -0.86 | 0.0021 |
| 1,3,9-Trimethyluric acid^III^ | 4.88 | 211.0895* | 0 | [M+H]^+^ |  | 196.0666, 183.0949, 167.0993, 154.0652, 139.0418, 126.0709, 124.0935, 111.0440, 110.0385, 83.0602 | Faecal pH | -3.08 | <0.0001 |
| Proline^I^ | 0.77 | 116.0715* | 3 | [M+H]^+^ |  | 70.0661* | Faecal pH | 0.36 | 0.0007 |

a) I-III: Level of identification: I, confirmed with an analytical standard; II, putatively annotated – based on a match to a spectral library; III, putatively annotated– based on spectral similarity to a chemical class^25^; b) FDR < 0.1 ; *Features selected by the two statistical models. Additional information can be found in **Supplementary Table 7**.

**Supplementary Table 7. Supporting information for the identification of metabolites is listed in Supplementary Table 6:**

| **4-Hydroxybenzoic acid-sulphate**, C_7_H_6_O_6_S  *m/z* = 216.9806 (-), RT = 3.89 | | |
| --- | --- | --- |
| 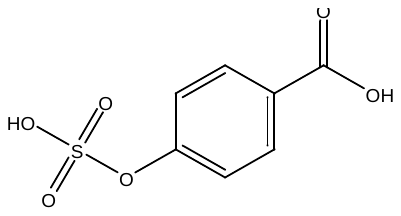 | [M-H]^-^ | 216.9806 |
|  | [M-SO_3_-H]^-^ | 137.0241 |
|  | [M- SO_3_-CO-H]^-^ | 109.0284 |
|  | [M- SO_3_-CO_2_-H]^-^ | 93.0344 |
| Identification level II; the compound matches with both 2-hydroxybenzoic acid-sulphate and 4-hydroxybenzoic acid-sulphate synthesized standards | | |

| **3-Hydroxy-2-oxindole sulphate**, C_8_H_7_NO_5_S,  *m/z*= 227.9967 (-), RT= 4.02 | | |
| --- | --- | --- |
| 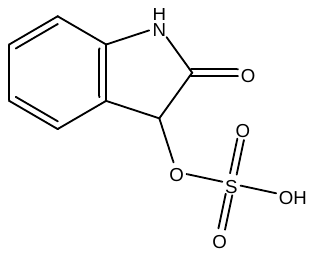 | [M-H]^-^ | 227.9967 |
|  |  | 186.0231 |
|  |  | 150.0559 |
|  | [M-SO_3_-H]^-^ | 148.0403 |
|  |  | 142.0329 |
|  | [M-SO_3_-CO-H]^-^  [M-SO_3_-CONH-H]^-^ | 120.0450  107.0380 |
|  | SO_3_^-^ | 79.9569 |
| Identification level III; putative annotation | | |

| **5-Hydroxy-2-oxindole sulphate**, C_8_H_6_NO_5_S,  *m/z*= 227.9967 (-), RT= 3.53 | | |
| --- | --- | --- |
| 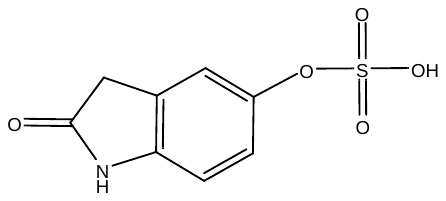 | [M-H]^-^ | 227.9967 |
|  |  | 182.9546 |
|  |  | 164.9429 |
|  |  | 151.0399 |
|  | [M-SO_3_]^-^ | 148.0399 |
|  |  | 145.098 |
|  |  | 138.9658 |
|  |  | 136.9493 |
|  | [M-SO_3_-CO-H]^-^ | 120.0451 |
|  | [M-SO_3_-CONH-H]^-^ | 107.0495 |
|  | [M-SO_3_-CONH_3_-H]^-^ | 105.0345 |
|  | [M-SO_3_-2CO-H]^-^ | 92.0502 |
|  | SO_3_^-^ | 79.9576 |
|  |  | 68.09 |
| Identification level II; the compound matches with the synthesized standard, but it is unknown whether the sulphate bonded to the hydroxy group in position 2 or 5. | | |

| **1-Methylxanthine**, C_6_H_6_N_4_O  *m/z*= 167.0555 (+) RT= 1.58 | | |
| --- | --- | --- |
| 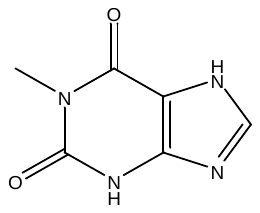 | [M+H]^+^ | 167.0555 |
|  | [M-C_3_H_5_O+H]^+^ | 110.0338 |
|  | [M-CONCH_3_-CO+H]^+^ | 82.0437 |
|  | [M-CONCH_3_-CO-HCN+H]^+^ | 55.0307 |
| Identification level I; matched to an authentic standard | | |

| **1,3-Dimethyluric acid**, C_6_H_6_N_4_O_3_  *m/z*= 195.0519 (-) RT= 2.00 | | |
| --- | --- | --- |
| 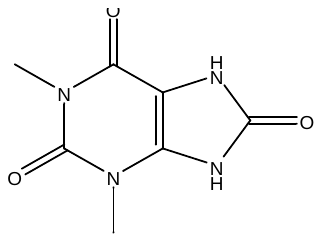 | [M-H]^-^ | 195.0519 |
|  | [M-CH_3_-H]^-^ | 180.0287 |
|  | [M-CH_3_NCO-H]^-^ | 138.0302 |
|  |  | 137.0227 |
|  | [M-CH_3_-CH_3_NCO-H]^-^ | 123.0075 |
|  | [M-CH_3_NCO-CO-H]^-^ | 110.0358 |
|  | [M-CH_3_NCO-CO-OH-H]^-^ | 93.0331 |
|  | [M-CH_3_NCO-CO-HCN-H]^-^ | 83.0253 |
|  | [M-CH_3_NCO-CO-CH_2_O-H]^-^ | 80.0257 |
| Identification level I; matched to an authentic standard | | |

| **1,7-Dimethyluric acid**, C_6_H_6_N_4_O_3_  *m/z*= 195.0526 (-) RT= 2.53 | | |
| --- | --- | --- |
| 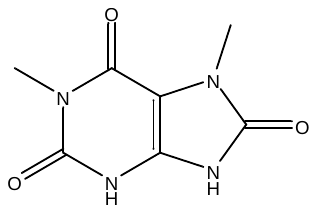 | [M-H]^-^ | 195.0526 |
|  | [M-CH_3_-H]^-^ | 180.0286 |
|  |  | 162.8392 |
|  |  | 160.8415 |
|  | [M-CH_3_NCO-H]^-^ | 138.0308 |
|  |  | 137.0228 |
|  | [M-CH_3_-CH_3_NCO-H]^-^ | 123.0067 |
|  |  | 121.9991 |
|  | [M-CH_3_NCO-CO-H]^-^ | 110.0355 |
|  | [M-CH_3_NCO-CO-OH-H]^-^ | 93.0350 |
|  | [M-CH_3_NCO-CO-OH-CN-H]^-^ | 67.0302 |
| Identification level II; matched to the spectral library (mzCloud) | | |

| **1,3,9-Trimethyluric acid**, C_8_H_10_N_4_O_3_  *m/z*= 211.0895 (+) RT= 2.79 | | |
| --- | --- | --- |
| 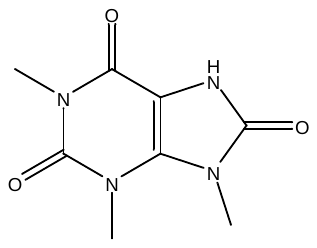 | [M+H]^+^ | 211.0895 |
|  | [M-CH_3_+H]^+^ | 196.0666 |
|  | [M-CO+H]^+^ | 183.0949 |
|  | [M-CO_2_+H]^+^ | 167.0993 |
|  | [M-CH_3_NCO+H]^+^ | 154.0652 |
|  | [M-CH_3_-CH_3_NCO+H]^+^ | 139.0418 |
|  | [M-CO-CH_3_NCO+H]^+^ | 126.0709 |
|  |  | 124.0935 |
|  | [M-CH_3_-CH_3_NCO-CO+H]^+^ | 111.0440 |
|  |  | 110.0385 |
|  | [M-CH_3_-CH_3_NCO-2CO+H]^+^ | 83.0602 |
| Identification level III; putative annotation | | |

| **Citric acid**, C_6_H_8_O_7_  *m/z*= 191.0192 (-) RT= 0.73 | | |
| --- | --- | --- |
| 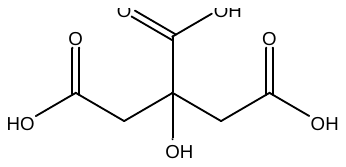 | [M-H]^-^ | 191.0192 |
|  | [M-H_2_O-H]^-^ | 173.0085 |
|  | [M-2H_2_O-H]^-^ | 154.9981 |
|  | [M-H_2_O-CO_2_-H]^-^ | 129.019 |
|  | [M-2H_2_O-CO_2_-H]^-^ | 111.0084 |
|  | [M-H_2_O-CO_2_-COCH_2_-H]^-^ | 87.0086 |
|  | [M-H_2_O-2CO_2_-H]^-^ | 85.0294 |
|  | [M-2H_2_O-2CO_2_-H]^-^ | 67.0188 |
|  | [M-H_2_O-2CO_2_-CO-H]^-^ | 57.0345 |
| Identification level I; matched to an authentic standard | | |

| **1-Methyluric acid**, C_6_H_6_N_4_O_2_  *m/z*= 181.0364 (-) RT= 1.31 | | |
| --- | --- | --- |
| 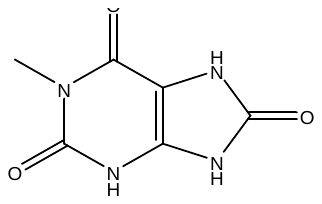 | [M-H]^-^ | 181.0364 |
|  | [M-CONH-H]^-^ | 138.0310 |
|  | [M-C_2_H_3_O-H]^-^ | 122.0230 |
|  | [M-CONH-CO-H]^-^ | 110.0350 |
|  | [M-CONH-COCH_2_-H]^-^ | 96.0203 |
|  | [M-CONH-CO-HCN-H]^-^ | 83.0244 |
|  | [M-CONH-COCH_2_-HCN-H]^-^ | 69.0091 |
| Identification level I; matched to an authentic standard | | |

| **Taurine**, C_2_H_7_NO_3_S,  *m/z*= 124.0065 (-) RT= 0.53 | | |
| --- | --- | --- |
| 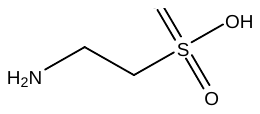 | [M-H]^-^ | 124.0065 |
|  | [M-NH_3_-H]^-^ | 106.9801 |
|  | [M-C_2_H_5_-H]^-^ | 94.9800 |
|  | [SO_3_]^-^ | 79.9569 |
| Identification level I; matched to an authentic standard | | |

| **4-Hydroxyhippuric acid-sulphate,** C_9_H_9_NO_7_S,  *m/z*= 274.0022 (-) RT= 3.86 | | |
| --- | --- | --- |
| 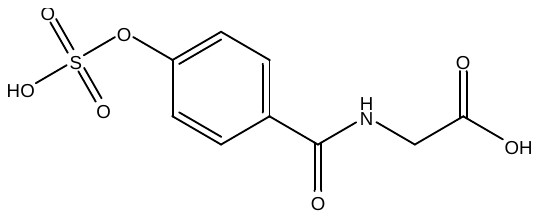 | [M-H]^-^ | 274.0022 |
|  | [M-CO_2_-H]^-^ | 230.0122 |
|  | [M-SO_3_-H]^-^ | 194.0458 |
|  | [M-SO_3_-CO_2_-H]^-^ | 150.0558 |
|  | [M-SO_3_-CO_2_-CH_2_NHCO-H]^-^ | 93.0354 |
| Identification level II;  the compound matches with both 4-hydroxyhippuric acid-sulphate and 3-hydroxyhippuric acid-sulphate synthesized standards | | |

| **Pseudouridine,** C_9_H_11_N_2_O_6_,  *m/z*= 243.0615 (-) RT= 0.65 | | |
| --- | --- | --- |
| 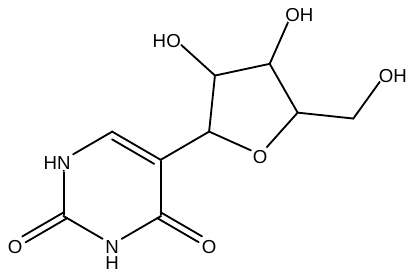 | [M-H]^-^ | 243.0615 |
|  | [M-H-H_2_O]^-^ | 225.0503 |
|  | [M-CH_2_O-CH_2_O]^-^ | 183.0405 |
|  | [M-CH_2_O-CH_2_O-CH_2_O]^-^ | 153.0297 |
|  | [M-CH_2_O-CH_2_O-CONH-H]^-^ | 140.0641 |
|  | [M-CH_2_O-CH_2_O-CH_2_O-CONH-H]^-^ | 110.0256 |
| Identification level I; matched to an authentic standard | | |

| **Pimelic acid**, C_7_H_12_O_4_  *m/z*= 159.0658 (-) RT= 3.23 | | |
| --- | --- | --- |
| 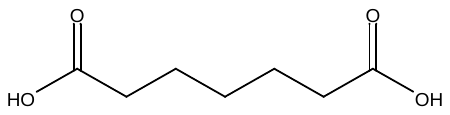 | [M-H]^-^ | 159.0658 |
|  | [M-H_2_O-H]^-^ | 141.0550 |
|  | [M-CO_2_-H]^-^ | 115.0761 |
|  | [M-H_2_O-CO_2_-H]^-^ | 97.0655 |
|  | [M-H_2_O-HCOOH-H]^-^ | 95.0489 |
| Identification level I; matched to an authentic standard | | |

| **4-Methylcatechol sulfate,** C_7_H_8_O_5_S  *m/z*= 203.0016 (-) RT= 3.42 | | |
| --- | --- | --- |
| 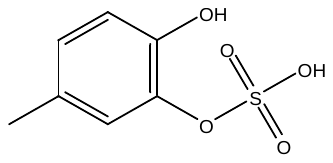 | [M-H]^-^ | 203.0016 |
|  | [M-SO_3_-H]^-^ | 123.0445 |
|  | [M-SO_3_]^.^ | 122.0369 |
|  | [M-SO_3_-CH_3_ -H]^-^ | 108.0216 |
|  | [M-SO_3_-OH -H]^-^ | 105.0343 |
|  | [M-SO_3_-CO -H]^-^ | 95.0505 |
|  | SO_3_- | 79.9871 |
| Identification level II; the compound matches with both 4-methylcatechol-sulphate and 3-methylcatechol-sulphate synthesized standards | | |

| **3-Hydroxy-2-oxindole glucuronide**, C_14_H_15_NO_8_,  *m/z*= 324.0724 (-) RT= 2.61 | | |
| --- | --- | --- |
| 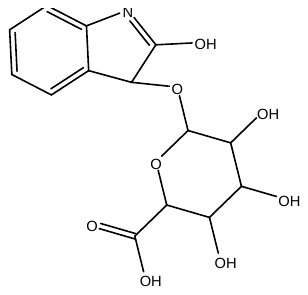 | [M-H]^-^ | 324.0724 |
|  | [C_6_H_10_O_7_-H]^-^ | 193.0351 |
|  | [C_6_H_10_O_7_-H_2_O-H]^-^ | 175.0246 |
|  |  | 160.0402 |
|  | [M-C_6_H_10_O_7_-H_2_O-H]^-^ | 148.0402 |
|  |  | 145.0143 |
|  | [C_6_H_10_O_7_-H_2_O-CO_2_-H]^-^ | 131.035 |
|  | [C_6_H_10_O_7_-2H_2_O-CO_2_-H]^-^ | 113.0242 |
|  |  | 103.0031 |
|  | [C_6_H_10_O_7_-H_2_O-CO_2_-CH_2_O-H]^-^ | 101.0244 |
|  |  | 89.0244 |
|  | [C_6_H_10_O_7_-2H_2_O-CO_2_-CO-H]^-^ | 85.0294 |
|  |  | 72.9924 |
|  | [C_6_H_10_O_7_-2H_2_O-CO_2_-CH_2_CO-H]^-^ | 71.0138 |
|  | [C_6_H_10_O_7_-2H_2_O-CO_2_-CO-C_2_H_2_-H]^-^ | 59.0134 |
| Identification level II; the compound matches with the synthesized standard, but it is unknown whether the sulphate bonded to the hydroxy group in position 2 or 3 | | |

| **Pipecolic acid**, C_6_H_11_NO_2_,  *m/z* = 130.0857 (+), RT = 0.99 | | |
| --- | --- | --- |
| 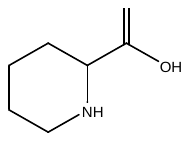 | [M+H]^+^ | 130.0857 |
|  | [M-H2O+H]^+^ | 112.0734 |
|  | [M-HCOOH+H]^+^ | 84.0802 |
|  | [C_3_H_5_N+H]+ | 56.0489 |
|  | [C_4_H_6_+H]+ | 55.0547 |
| Identification level I, matched to an authentic standard | | |

| **Glutaric acid**, C_5_H_8_O_4_,  *m/z* = 131.0359 (-), RT = 3.20 | | |
| --- | --- | --- |
| 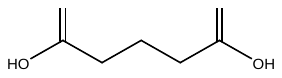 | [M-H]^-^ | 131.0359 |
|  | [M-CO_2_-H]^-^ | 87.04452 |
|  | [M-HCOOH-H]^-^ | 85.0295 |
|  |  | 76.98276 |
|  | [M-CO_2_-H_2_O-H]^-^ | 69.0351 |
|  | [CH_3_COOH-H]^-^ | 59.0142 |
| Identification level I; matched to an authentic standard | | |

| **Suberic acid**, C_8_H_14_O_4_,  *m/z* = 173.0814 (-), RT = 5.72 | | |
| --- | --- | --- |
| 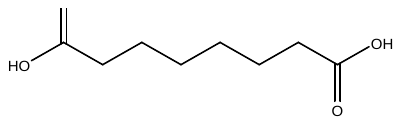 | [M-H]^-^ | 173.0814 |
|  | [M-CO_2_-H]^-^ | 129.0912 |
|  | [M-CO_2_-H_2_O-H]^-^ | 111.0814 |
|  |  | 102.9716 |
|  |  | 86.9762 |
|  | [M-CO_2_-H_2_O-C_2_H_4_-H]^-^ | 83.0504 |
| Identification level I; matched to an authentic standard | | |

| **Sebacic acid**, C_10_H_18_O_4_,  *m/z* = 201.1123 (-), RT = 6.29 | | |
| --- | --- | --- |
| 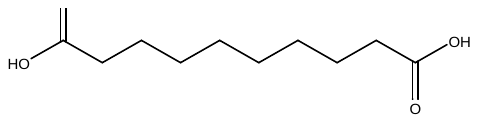 | [M-H]^-^ | 201.1123 |
|  | [M-OH-H]^-^ | 183.1020 |
|  | [M-CO_2_-H_2_O-H]^-^ | 139.1124 |
|  | [M-H_2_O-CO_2_-C_2_H_4_-H]^-^ | 111.0826 |
|  |  | 61.9879 |
| Identification level I; matched to an authentic standard | | |

| **Xanthine**, C_5_H_4_N_4_O_2_,  *m/z* = 151.0254 (-), RT = 1.66 | | |
| --- | --- | --- |
| 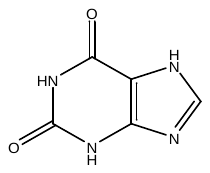 | [M-H]^-^ | 151.0254 |
|  | [M-H_2_O-H]^-^ | 133.0147 |
|  | [M-CH_3_CO-H]^-^ | 108.0197 |
|  | [M-CH_3_CO-CO-H]^-^ | 80.0263 |
|  | [M-CONH-COCH_2_-H]^-^ | 65.9971 |
| Identification level I; matched to an authentic standard | | |

| **Deoxyxanthosine**, C_11_H_13_N_4_O_4_,  *m/z* = 267.0769 (-), RT = 4.04 | | |
| --- | --- | --- |
| 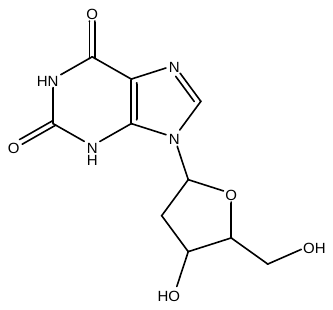 | [M-H]^-^ | 267.0769 |
|  | [M-C_3_H_6_O-H]^-^ | 177.0404 |
|  | [M-C_5_H_10_O_4_-H]^-^ | 151.0259 |
|  | [M-C_3_H_6_O-CONH-H]^-^ | 134.0360 |
|  | [M- C_5_H_10_O_4_-CONH-H]^-^ | 108.0200 |
|  | [M- C_5_H_10_O_4_-CONH-COCH_2_-H]^-^ | 92.0254 |
| Identification level III; putative annotation | | |

| **Tryptophan**, C_11_H_12_N_2_O_2_,  *m/z* = 205.0983 (+), RT = 4.44 | | |
| --- | --- | --- |
| 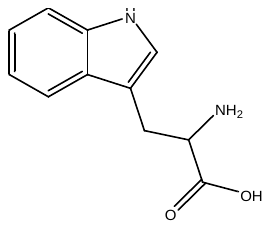 | [M+H]^+^ | 205.0983 |
|  | [M-NH_2_+H]^+^ | 188.0698 |
|  | [M-NH_2_-H_2_O+H]^+^ | 170.0592 |
|  | [M-HCOOH+H]^+^ | 159.0906 |
|  | [M-NH_2_-COCH_2_+H]^+^ | 146.0593 |
|  | [M-NH_2_-CO_2_+H]^+^ | 144.0797 |
|  | [M-HCOOH-HCN+H]^+^ | 132.08155 |
|  | [M-NH_2_-COCH_2_-CO+H]^+^ | 118.0649 |
| Identification level I; matched to an authentic standard | | |

| **2-Oxindole-3-acetic acid**, C_9_H_7_NO_3_,  *m/z* = 192.0655 (+), RT = 5.36 | | |
| --- | --- | --- |
| 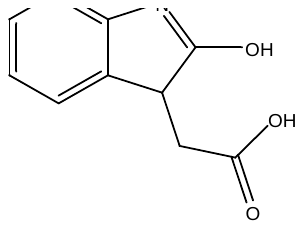 | [M+H]^+^ | 192.0655 |
|  | [M-H_2_O+H]^+^ | 174.0544 |
|  | [M-HCOOH+H]^+^ | 146.0596 |
|  |  | 135.0433 |
|  | [M-HCOOH-H_2_O+H]^+^ | 128.0493 |
|  |  | 120.0800 |
|  | [M-HCOOH-CO+H]^+^ | 118.0648 |
|  |  | 107.0748 |
|  | [M-HCOOH-COCH_2_+H]^+^ | 104.0491 |
|  | [M-HCOOH-H_2_O-HCN+H]^+^ | 101.0378 |
|  | [M-HCOOH-CO-HCN+H]^+^ | 91.0552 |
|  |  | 79.0554 |
|  | [M-HCOOH-COCH_2_-HCN+H]^+^ | 77.0385 |
| Identification level I; matched with authentic standard | | |

| **Pantothenic acid**, C_9_H_17_NO_5_,  *m/z* = 220.1172(+), RT = 4.19 | | |
| --- | --- | --- |
| 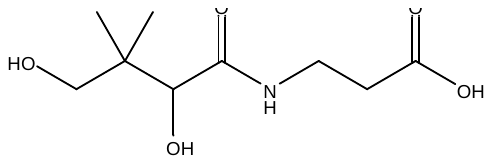 | [M+H]^+^ | 220.1172 |
|  | [M-H_2_O+H]^+^ | 202.1067 |
|  | [M-2H_2_O +H]^+^ | 184.0962 |
|  | [M-2H_2_O-COCH_2_+H]^+^ | 142.0847 |
|  | [M-3H_2_O-COCH_2_+H]^+^ | 124.075 |
|  | [M-2H_2_O-C_5_H_8_+H]^+^ | 116.0333 |
|  | [M-3H_2_O-C_5_H_8_+H]^+^ | 98.0236 |
|  | [C_3_H_7_O_2_N+H]^+^ | 90.0544 |
|  | [C_3_H_5_ON+H]^+^ | 72.0432 |
|  | [M-2H_2_O-COCH_2_+H]^+^ | 67.055 |
| Identification level I; matched to an authentic standard | | |

| **Nicotinic acid**, C_6_H_5_NO_2_,  *m/z* = 124.0402 (+), RT = 1.17 | | |
| --- | --- | --- |
| 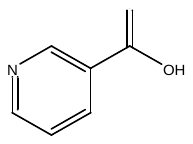 | [M+H]^+^ | 124.04 |
|  | [M-H_2_O+H]^+^ | 106.029 |
|  | [M-CO+H]^+^ | 96.0451 |
|  | [M-CO_2_+H]^+^ | 80.0493 |
|  | [M-HCOOH+H]^+^ | 78.0332 |
| Identification level I; matched to an authentic standard | | |

| **Proline**, C_5_H_9_NO_2_,  *m/z* = 116.0699 (+), RT = 0.77 | | |
| --- | --- | --- |
| 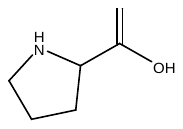 | [M+H]^+^ | 116.0699 |
|  | [M+HCOOH+H]^+^ | 70.0646 |
| Identification level I; matched to an authentic standard | | |

| **4-Hydroxyphenyllactic acid**, C_9_H_10_O_4_,  *m/z* = 181.0495 (-), RT = 2.62 | | |
| --- | --- | --- |
| 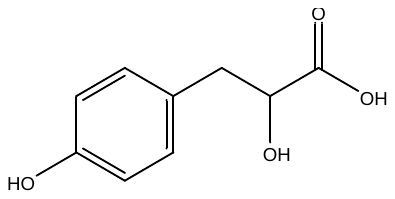 | [M-H]^-^ | 181.0495 |
|  | [M-H_2_O-H]^-^ | 163.0398 |
|  | [M-HCOOH-H]^-^ | 135.044 |
|  | [M-H_2_O-CO_2_]^-^ | 119.0493 |
|  | [M-HCOOH-CO-H]^-^ | 107.0496 |
|  | [(CHOH-COOH)-H]^-^ | 72.9947 |
| Identification level II; matched to an authentic standard | | |

| **Picolinoylglycine**, C_8_H_8_N_2_O_3_,  *m/z*= 181.0702 (+), RT= 4.66 | | |
| --- | --- | --- |
| 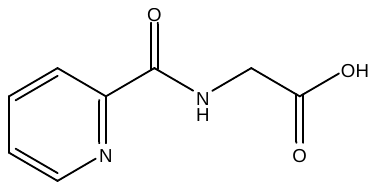 | [M+H]^+^ | 181.0702 |
|  | [M-H_2_O+H]^+^ | 163.0485 |
|  | [M-HCOOH+H]^+^ | 135.0544 |
|  | [M-NHCH_2_CO+H]^+^ | 124.0489 |
|  | [M-Gly+H]^+^ | 106.0276 |
|  |  | 96.0431 |
|  | [M-Gly-CO+H]^+^ | 78.0325 |
|  |  | 69.0439 |
|  |  | 67.0281 |
|  |  | 55.0284 |
|  | [C_4_H_2_+H]^+^ | 51.0199 |
| Identification level III; putative annotation | | |

| **Dihydroferulic acid glucuronide**, C_16_H_20_O_10_,  *m/z*= 371.0981 (-), RT= 4.97 | | |
| --- | --- | --- |
| 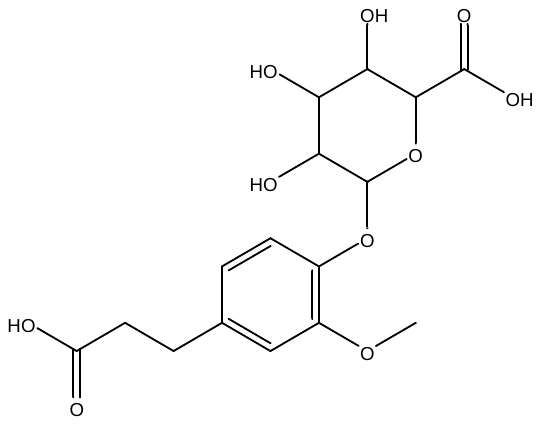 | [M-H]^-^ | 371.0981 |
|  | [M-H_2_O-H]^-^ | 353.0882 |
|  |  | 326.1336 |
|  | M-H_2_O-CO_2_-H]^-^ | 291.0848 |
|  | [M-(C_6_H_10_O_7_-H_2_O)-H]^-^ | 195.066 |
|  | [C_6_H_10_O_7_-H_2_O-H]^-^ | 175.0246 |
|  | [M-(C_6_H_10_O_7_-H2O)-CH_3_COOH-H]^-^ | 135.0445 |
|  | [C_6_H_10_O_7_-2H_2_O-CO_2_-H]^-^ | 113.0244 |
|  | [C_6_H_10_O_7_-2H_2_O-CO_2_-CO-H]^-^ | 85.0293 |
|  | [C_6_H_10_O_7_-2H_2_O-CO_2_-CH_2_CO-H]^-^ | 71.0128 |
|  | [C_6_H_10_O_7_-2H_2_O-CO_2_-CO-C_2_H_2_-H]^-^ | 59.0145 |
| Identification level III; putative annotation | | |

| **Argininic acid**, C_6_H_13_N_3_O_3_,  *m/z*= 176.1020 (+), RT= 0.93 | | |
| --- | --- | --- |
| 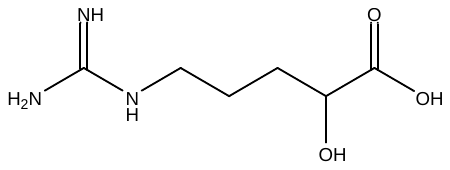 | [M+H]^+^ | 176.1020 |
|  | [M-NH_3_+H]^+^ | 159.0773 |
|  | [M-CH_2_N_2_+H]^+^ | 134.0802 |
|  | [M-HCOOH+H]^+^ | 130.0963 |
|  | [C_6_H_9_NO+H]^+^ | 112.0751 |
|  | [C_6_H_9_NO-CO+H]^+^ | 84.0810 |
|  | [C_6_H_9_NO-CH_5_N_3_+H | 71.0492 |
|  | [M-HCOOH-H_2_O-CNHNH_2_+H]^+^ | 70.0674 |
|  | [CH_5_N_3_+H]^+^ | 60.0575 |
| Identification level III; putative annotation | | |

| **Ascorbic acid glucuronide,** C_12_H_16_O_12_  *m/z*= 351.0569 (-), RT= 1.02 | | |
| --- | --- | --- |
| 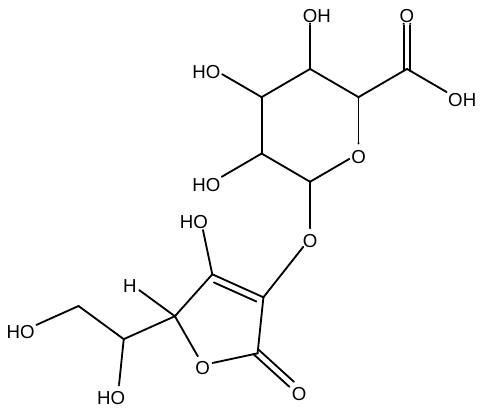 | [M-H]^-^ | 351.0569 |
|  | [M-H_2_O-H]^-^ | 333.0477 |
|  | [M-H_2_O-CO_2_-H]^-^ | 291.0357 |
|  |  | 241.9762 |
|  |  | 235.0393 |
|  | [C_6_H_10_O_7_-H]^-^ | 193.0348 |
|  | [M-(C_6_H_10_O_7_-H_2_O)-H]^-^ | 175.0246 |
|  |  | 162.0196 |
|  | [M-C_6_H_10_O_7_-H_2_O-H]^-^ | 157.0140 |
|  | [M-C_6_H_10_O_7_-H_2_O-CO_2_-H]^-^ | 115.0035 |
|  | [M-C_6_H_10_O_7_-H_2_O-CO_2_-CO-H]^-^ | 87.0087 |
| Identification level III; putative annotation | | |

| **Unknown bile acid glycine conjugate**  *m/z* = 470.3422 (+), RT = 6.78 | | | |
| --- | --- | --- | --- |
|  | | [M+H]^+^ | 470.3422 |
|  |  | [M+H-H_2_O]^+^ | 452.3318 |
|  |  | [M+H-2H_2_O]^+^ | 434.3215 |
|  |  | [M+H-3H_2_O]^+^ | 416.3105 |
|  |  | [M+H-Gly]^+^ | 341.2781 |
|  |  | [M+H-Gly-H_2_O]^+^ | 323.2611 |
|  |  |  | 297.2496 |
|  |  |  | 227.1424 |
|  |  |  | 209.1315 |
|  |  |  | 171.1384 |
|  |  |  | 158.0811 |
|  |  |  | 123.0807 |
|  |  |  | 110.1054 |
|  |  |  | 109.0653 |
|  |  |  | 107.0845 |
|  |  |  | 95.0856 |
|  |  |  | 93.0686 |
|  |  |  | 76.0404 |
|  |  |  | 69.0731 |
| Identification level III; putative annotation | | | |

| **Cholic acid**, C_24_H_40_O_5_,  *m/z* = 407.2796 (-), RT = 6.74 | | |
| --- | --- | --- |
|  | [M-H]^-^ | 407.2796 |
| 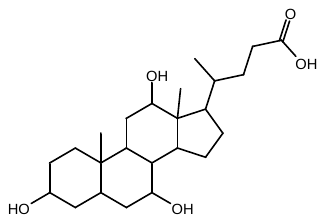 |  | 390.9812 |
|  |  | 386.9874 |
|  |  | 366.9807 |
|  |  | 350.9871 |
|  |  | 345.2803 |
|  |  | 342.9791 |
|  |  | 340.9842 |
|  |  | 324.9888 |
|  |  | 304.9824 |
|  |  | 302.986 |
|  |  | 290.9892 |
|  |  | 270.9793 |
|  |  | 264.9896 |
|  |  | 254.9873 |
|  |  | 242.9851 |
|  |  | 240.9892 |
|  |  | 204.9896 |
|  |  | 192.9872 |
|  |  | 191.943 |
|  |  | 174.9574 |
|  |  | 154.9916 |
|  |  | 146.9877 |
|  |  | 130.9921 |
|  |  | 92.566 |
| Identification level III; putative annotation | | |

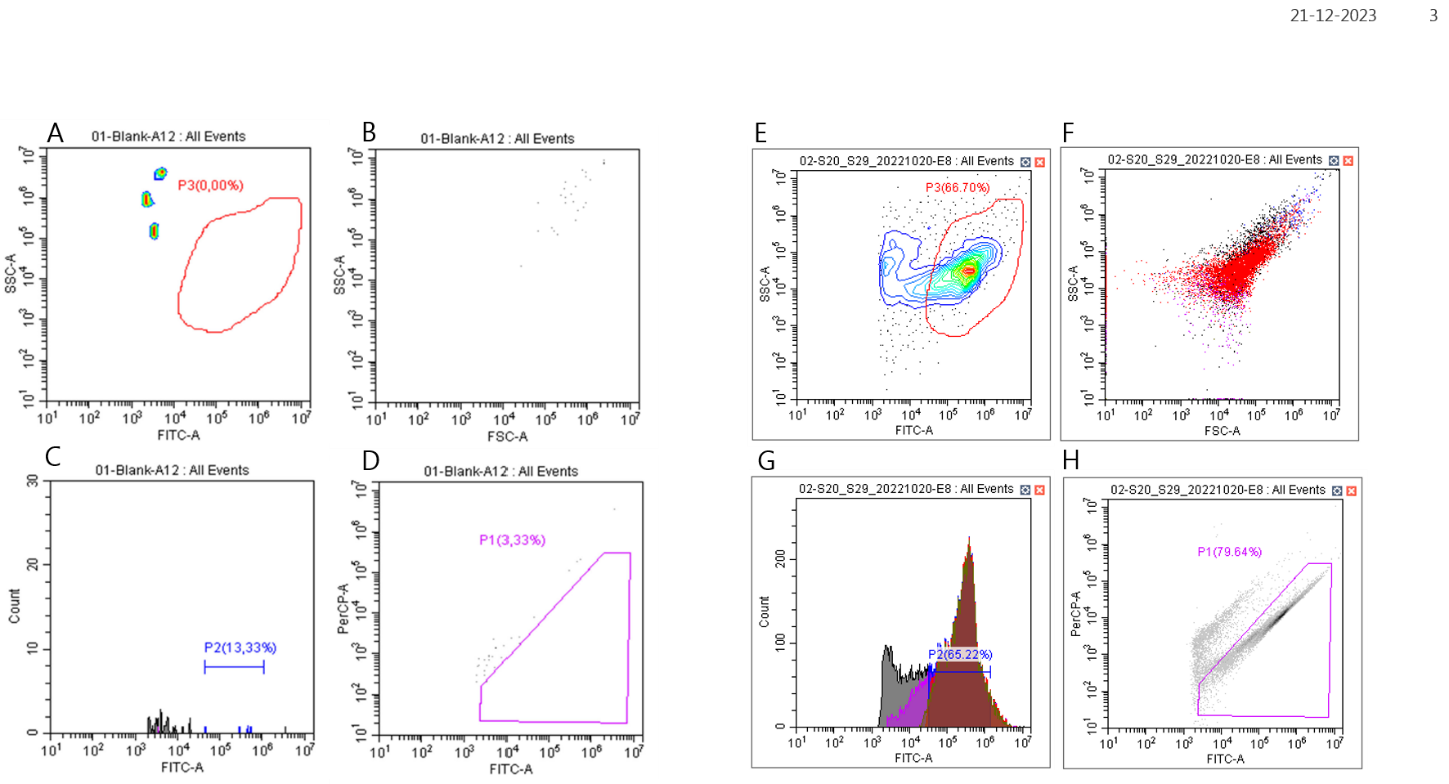

**Supplementary Figure 8.** A fixed flow cytometry gating and staining approach was applied. Both blank and sample solutions were stained with SYBR Green I. Fluorescence events were measured using fluorescence channels FITC 525/40 nm combining with Side Scatter Channel SSC-A (A and E). In addition, backward gating at FSC-A combining SSC-A dot plot (B and F), FICT histogram plot (C and G) and FITC combining PerCP dot plot (D and H) was applied to double check the distribution of detected cells. The measured cells have similar size and fluorescence density, have a clear peak with normal distribution.

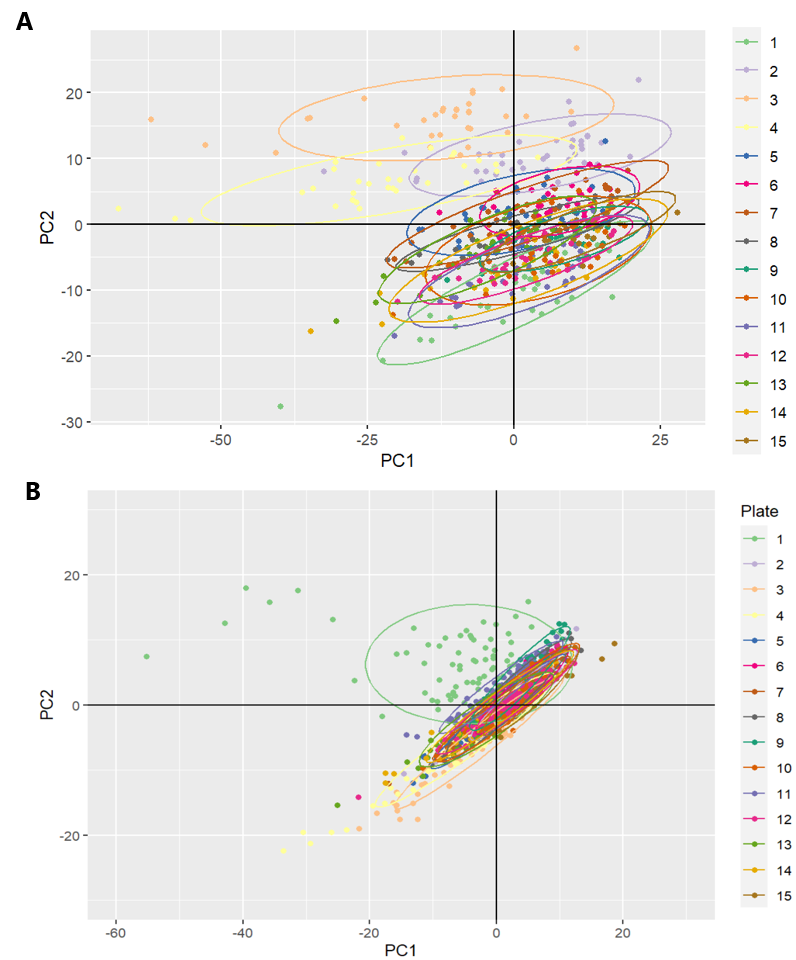

**Supplementary Figure 9.** PCA scores plot of QC samples detected in urine in A) negative mode and B) positive mode on LC-MS coloured by the plate. Plate no. 3 and 4 in negative mode and plate no. 1 in positive mode were excluded from further data analysis as they were clear outliers.

**Supplementary Table 8. LC-MS pre-processing parameters**

| **Faecal metabolomics** | | |
| --- | --- | --- |
| **Ionization mode** | **POS** | **NEG** |
| **CentWave algorithm prefilter** | scans = 3, intensity = 10 | scans = 3, intensity = 10 |
| **Signal-to-noise threshold** | 100 | 100 |
| **Maximum m/z deviation tolerance** | 50 | 50 |
| **Peak width** | 0.015-0.2 | 0.0125-0.2 |
| **Integrate** | 2 | 2 |
| **Gap filling** | expandMz = 0, expandRt = 0, ppm = 30) | expandMz = 0, expandRt = 0, ppm = 30) |
| **Urine metabolomics** | | |
| **Ionization mode** | **POS** | **NEG** |
| **CentWave algorithm prefilter** | scans = 3, intensity = 10 | scans = 3, intensity = 10 |
| **Signal-to-noise threshold** | 10 | 10 |
| **Maximum m/z deviation tolerance** | 30 | 30 |
| **Peak width** | 0.01-0.2s | 0.01-0.2s |
| **Integrate** | 2 | 2 |
| **Gap filling** | expandMz = 0, expandRt = 0, ppm = 30) | expandMz = 0, expandRt = 0, ppm = 30) |
